# Hybrid Automata Library: A modular platform for efficient hybrid modeling with real-time visualization

**DOI:** 10.1101/411538

**Authors:** Rafael Bravo, Etienne Baratchart, Jeffrey West, Ryan O. Schenck, Anna K. Miller, Jill Gallaher, Chandler D. Gatenbee, David Basanta, Mark Robertson-Tessi, Alexander R. A. Anderson

## Abstract

The Hybrid Automata Library (HAL) is a Java Library developed for use in mathematical oncology modeling. It is made of simple, efficient, generic components that can be used to model complex spatial systems. HAL’s components can broadly be classified into: on- and off-lattice agent containers, finite difference diffusion fields, a GUI building system, and additional tools and utilities for computation and data collection. These components are designed to operate independently and are standardized to make them easy to interface with one another. As a demonstration of how modeling can be simplified using our approach, we have included a complete example of a hybrid model (a spatial model with interacting agent-based and PDE components). HAL is a useful asset for researchers who wish to build efficient 1D, 2D and 3D hybrid models in Java, while not starting entirely from scratch. It is available on github at https://github.com/MathOnco/HAL under the MIT License. HAL requires at least Java 8 or later to run, and the Java JDK version 1.8 or later to compile the source code.

**Author Summary:** In this paper we introduce the Hybrid Automata Library (HAL) with the purpose of simplifying the implementation and sharing of hybrid models for use in mathematical oncology. Hybrid modeling is used in oncology to create spatial models of tissue, typically by modeling cells using agent-based techniques, and by modeling diffusible chemicals using partial differential equations (PDEs). HAL’s key components are designed to run agent-based models, PDEs, and visualization. The components are standardized and are completely decoupled, so models can be built with any combination of them. We first explore the philosophy behind HAL, then summarize the components. Lastly we demonstrate how the components work together with an example of a hybrid model, and a walk-through of the code used to construct it. HAL is open-source and will produce identical results on any machine that supports Java 8 and above, making it highly portable. We recommend HAL to modelers interested in spatial dynamics, even those outside of mathematical oncology, as the components are general enough to facilitate a variety of model types. A community page that provides a download link and online documentation can be found at https://halloworld.org [1].

## 2 Introduction

The Hybrid Automata Library (HAL) was created to assist the growing mathematical oncology community with a common framework for efficiently building and visualizing hybrid models. Hybrid models in oncology usually represent cells (both of the tumor and of the surrounding tissue) using agent-based modeling (ABMs) and the concentrations of relevant chemicals (drugs, resources and signaling molecules) as continuous partial differential equations (PDEs). These models can simulate local interactions between cells with complex internal dynamics and decision-making processes while also allowing cells to interact with the PDE concentration fields in their local environment.

Hybrid models have been widely adopted within the Mathematical Oncology community to model many aspects of cancer [2–5]. A unique strength of the hybrid modeling approach is that it allows for a mechanistic understanding of the ecological feedback between the cancer cells and their tissue environment. Cancer cell agents can be modeled as a part of the surrounding tissue, and interact with the systems that normally maintain homeostasis. [6–11] Drugs may be subsequently introduced to add additional selective pressure to the model, and the long-term effects on the tumor evolution observed. A better understanding of these selection dynamics can be used to help develop more effective drug sequences to prevent cancer resistance to therapy and to develop evolutionary therapies to control cancers that cannot be cured with maximum tolerated dose [12–14]. Further realism can be incorporated by initializing spatial models with clinical or histological data [15, 16].

Whilst a number of agent-based modeling frameworks have been used for tissue modeling, including MASON, Repast, Physicell, CompuCell3D, Chaste, and Netlogo, we designed HAL to be simpler, more efficient, and more flexible.

Some of these frameworks facilitate model building under specific spatial interaction assumptions like PhysiCell [17], which treats cells as spheres under Newtonian adhesion-repulsion dynamics and is optimized for large cell populations, and CompuCell 3D [18], which models cells as contiguous composites of lattice positions, allowing cell deformation. HAL does not include the same depth in the domains specific to these frameworks, but uses a broader approach to provide the capacity for a variety of approaches.

Some of the most popular frameworks that also take a broad approach are Chaste, Repast, Mason, and Netlogo. Chaste uses an assumption based system for model building, in which modular rules are composed to define behavior, and behaviors that are not currently represented can be added as new modules [19]. This modular approach allows for very rapid prototyping, and increases the reproducibility of results. Repast uses a hierarchical nesting approach to group agents into sets that will all execute some action, and also features a highly customizable scheduling procedure to sequence these actions [20]. MASON is probably the most architecturally similar to HAL, as it also strives to be a modular agent-based modeling package, with built-in optional visualization tools and comparatively lax structure [21]. Netlogo uses a custom scripting language in order to simplify the coding process [22]. Netlogo also provides an accessible model development environment, making it a great choice for new modelers/coders. Each of these frameworks facilitates modeling under a different centralized scheduling structure, as mentioned in Table 1.

**Table 1.**
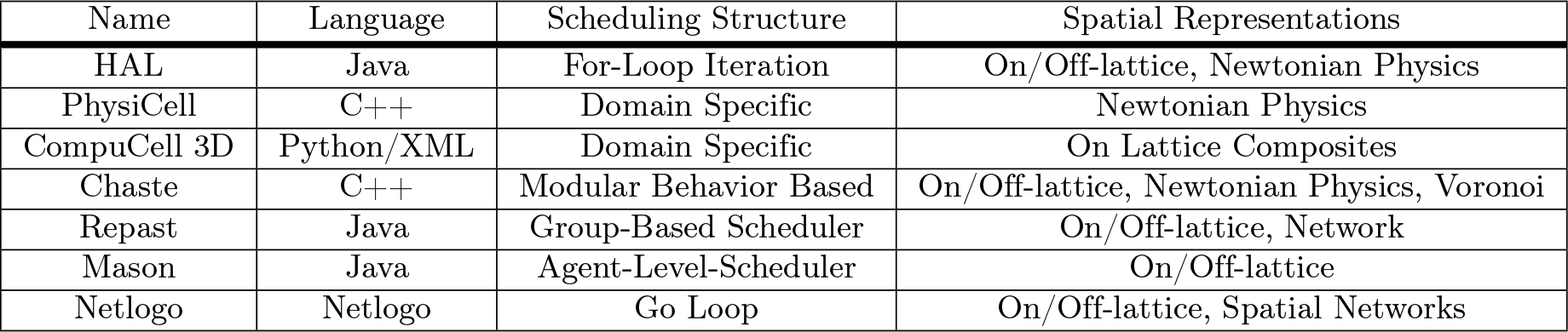
Comparison of HAL with other agent-based Modeling Frameworks commonly used in tissue modeling

| Name | Language | Scheduling Structure | Spatial Representations |
| --- | --- | --- | --- |
| HAL | Java | For-Loop Iteration | On/Off-lattice, Newtonian Physics |
| PhysiCell | C++ | Domain Specific | Newtonian Physics |
| CompuCell 3D | Python/XML | Domain Specific | On Lattice Composites |
| Chaste | C++ | Modular Behavior Based | On/Off-lattice, Newtonian Physics, Voronoi |
| Repast | Java | Group-Based Scheduler | On/Off-lattice, Network |
| Mason | Java | Agent-Level-Scheduler | On/Off-lattice |
| Netlogo | Netlogo | Go Loop | On/Off-lattice, Spatial Networks |

HAL shares many characteristics with these frameworks, but differentiates itself with a minimal, decentralized design made up of independent building blocks that are thematically similar. There is no centralized controller or scheduler, so the modeler designs the logical flow and the scheduling of interactions between model components. This removes common presuppositions or requirements made by schedulers in other frameworks (eg. when models should be visualized, when their step logic should run, when models should be created or destroyed, etc.) and leaves these decisions up to the modeler. This cuts down on any unnecessary use of resources by the modeling system, and increases model flexibility. These considerations have led to a lightweight framework that is easy to use, highly flexible, and effective within the scope of hybrid modeling, agent-based modeling, and the solving of reaction-diffusion PDEs using finite differences. HAL was designed with mathematical oncology in mind, but is general enough to facilitate modeling systems from many domains (eg. ecology [23], development, population dynamics, and network theory). [17]. While some familiarity with the Java programming language is recommended for new users of HAL, we imagine that its simplicity and explicit nature could make it a useful educational platform.

The main components of HAL consist of n-dimensional (0D,1D,2D,3D) Grids that hold Agents, 1D,2D, and 3D finite difference PDE fields, 2D and 3D visualization tools, and methods for sampling distributions and data recording. In this paper we will discuss the philosophy behind these components, then look at their design and capabilities in more detail. See the manual for a complete reference on how to use these components [24].

## 3 Design And Implementation

### 3.1 Design Philosophy

In the next section, we discuss some of the design decisions that have driven the architecture of HAL.

### 3.1.1 Language Choice

In designing HAL we have tried to balance an adherence to speed, memory management, simplicity, stability, modularity and extensibility. The Java language itself balances these considerations very well, making it a suitable basis for HAL. High performance languages such as C, C++, and Fortran, can be coded to run at speeds comparable to or faster than Java, however these languages require more low-level management. Moreover, they do not have the same security guarantees as they permit out-of-bounds memory accesses and memory leaks. Higher level languages, such as Python, while more flexible and syntactically intuitive than Java, are typically significantly slower. Java is also one of the most commonly used and taught programming languages today, which helps facilitate the adoption of HAL by new users. The fact that Java is cross-platform is also ideal.

#### 3.1.2 Modularity and Extensibility

HAL’s components can each function independently. This permits any number of components to be used in a single model, with the use of spatial queries to combine components, as seen in Fig 1. This modularity also allows modelers to choose only the components of HAL that are of interest for their project. These components can be easily mixed and matched with other software, such as using the AgentGrids with a different PDE solver, or using the GUI and Visualization components with a different modeling system. Modularity also makes adding new components more manageable and easier to test without adding bulk or heavy modifications to the core of the platform.

**Figure 1.**
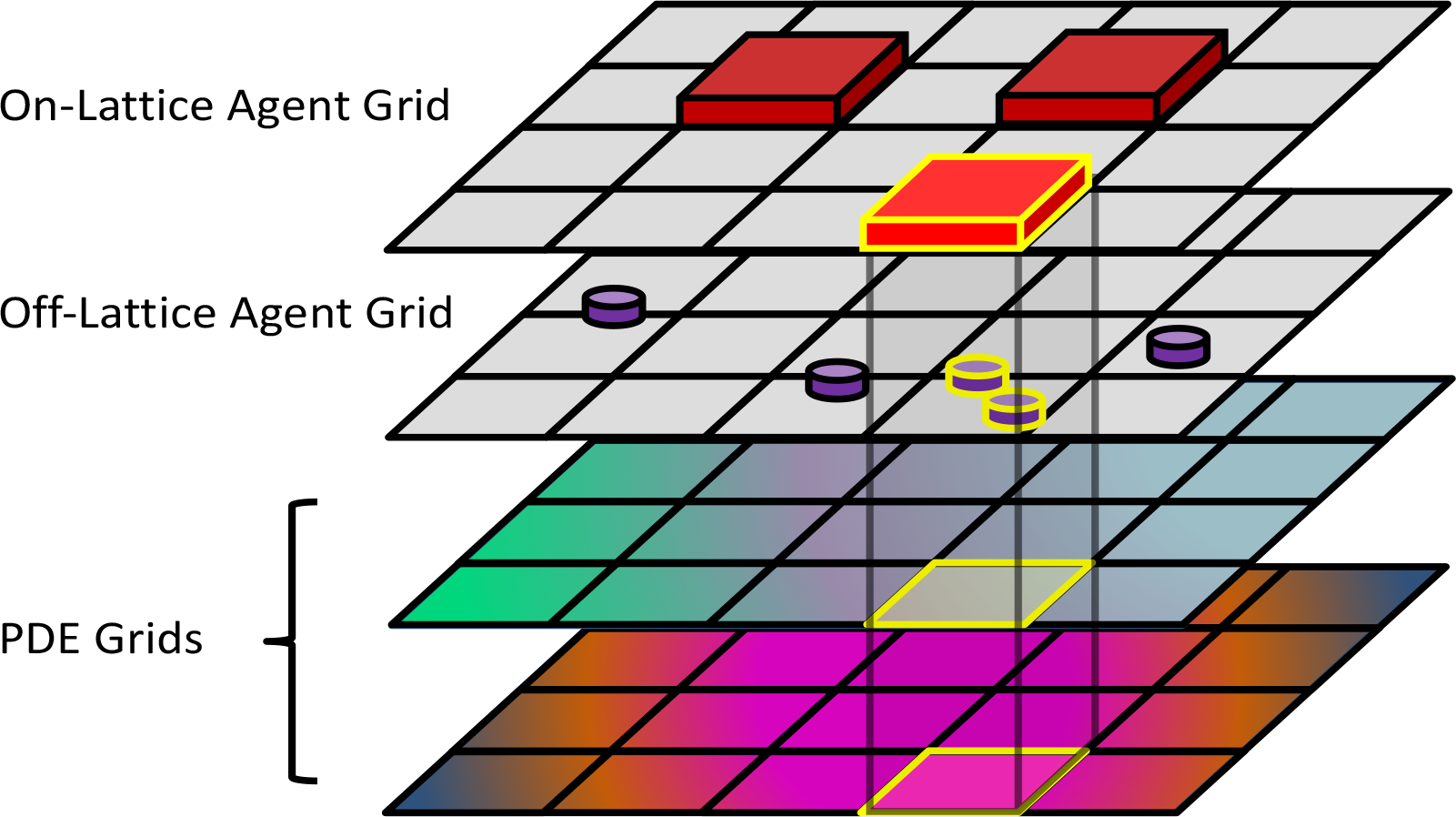
The modular design of HAL helps build complex models out of simple components. The highlighted on-lattice agent in the topmost grid searches for local overlaps with several other grids and PDEs. These overlaps can be used in a model to generate spatial interactions.

Given the incremental nature of many scientific endeavors, we also wanted to allow models and components to be extended and modified. Java’s extension architecture provides an excellent environment for layered development. As an example of the extensibility of HAL, the built-in Spherical Agent types (SphericalAgent2D, SphericalAgent3D) extend the Point Agent types (AgentPT2D, AgentPT3D). By default, Point Agents have no radius and will not collide with each other. This behavior can be useful for modeling phenomena such as the Brownian motion of small particles, as visualized in Fig 2a. Spherical Agents extend Point Agents by adding an additional radius variable and velocity component variables. These properties combined with added functions for summing force vectors in response to overlap allow for a Newtonian adhesion-repulsion spherical model of spatial agents. This behavior can be useful for modeling tissue formation, as visualized in Fig 2b.

**Figure 2.**
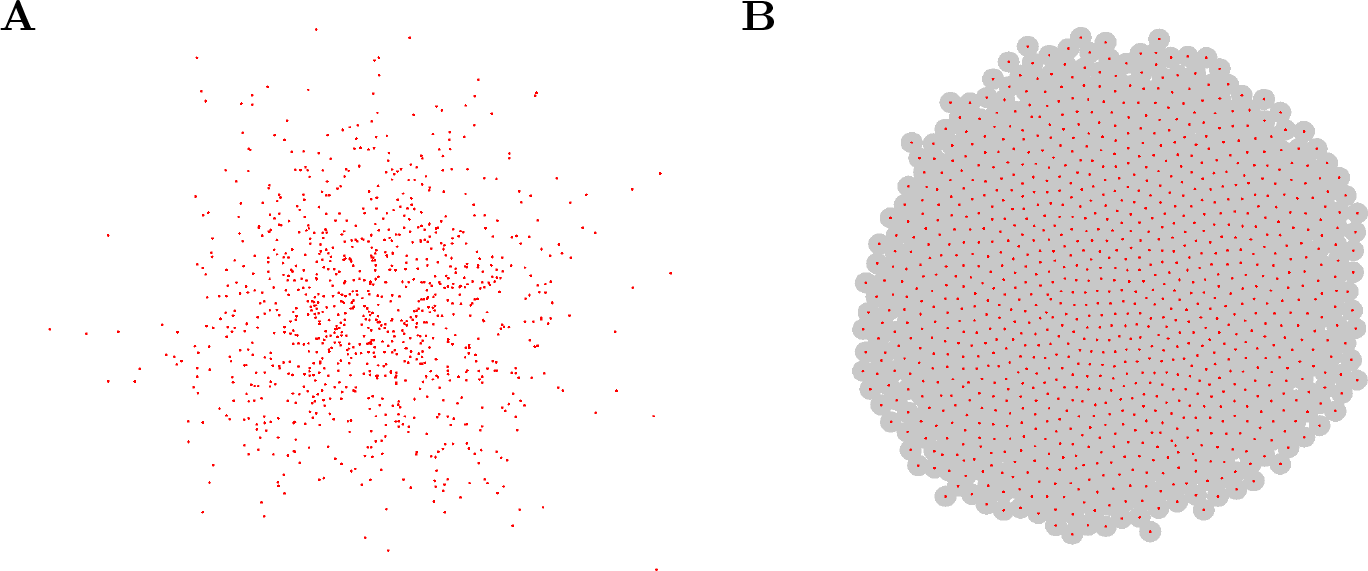
Off-lattice agent examples. Each example contains 1000 agents. **(A)** Example of 2D Point Agents modeling Brownian motion. The Point Agents move freely and cannot collide. **(B)** Example of 2D Spherical Agents modeling growing tissue. The agents will push apart from each other to a uniform density. Agent radii are shown as gray circles around their centers. Examples Displayed unsing the OpenGL2DWindow object.

It is also possible to extend completed models using the same approach. For example, grids and agents from published models can be used as a scaffold on which to do additional studies. This allows for followup studies to focus on implementing whatever additional assumptions and functionality they need, while leaving intact the base model code with all of its published assumptions.

#### 3.1.3 Simplicity and Stability

An important design principle was to make HAL simple to use without sacrificing performance. Simplicity makes HAL easy to learn and forces the components to be more generic, meaning that the same components can be applied to a greater variety of modeling problems. There is also a consistency to each framework component, such that learning to use some components is often sufficient to grasp the others, and makes using them in combination intuitive.

Another key design principle is stability, which is achieved in three ways:

Encapsulation: By providing safe interaction functions and preventing direct interaction with component internals. For example, modelers are not permitted to directly modify the position properties of agents. Instead, they must call the provided movement functions that also update the grid position of the agents for future spatial queries.
Defensive Programming: By including checks in functions for invalid inputs. The program halts and throws an error message immediately when one of these problematic inputs occurs. This allows the user to see what caused the problem, rather than seeing its effects later down the line.
Unit Tests: By testing HAL’s components. HAL is very shallow by design, leaving little complexity for bugs to hide in. The more complex algorithms are tested in a series of small test programs. These tests help ensure confidence in the mathematics while also serving as simple tutorials.

#### 3.1.4 Speed and Memory Management

Much of the performance capability of HAL comes directly from its decentralized design. Having no built-in scheduler/underlying structure means that there is comparatively little work that the program does that the modeler is unaware of. This combined with the modular components and utilities allows modelers the flexibility to incorporate the functionality that they need, without the software sacrificing performance by implicitly doing unnecessary tasks.

HAL also prioritizes performance in its algorithmic implementation. HAL includes efficient PDE solving algorithms, such as the ADI (alternating direction implicit) method, and uses efficient distribution samplers rather than naive approaches. The integrated visualization tools are also highly efficient, using Swing BufferedImages for lattice-based visualization, and lwjgl OpenGL for 2D and 3D polygon graphics. Whenever possible, primitives and arrays are used to store data rather than classes, which takes advantage of Java’s optimization for these simpler data types. Java is also an inherently fast language, which helps efficiently execute agent behavioral logic.

There is a memory footprint consideration with most of HAL’s assets. A common criticism of Java applications is that they tend to use a lot of memory and are slowed down by Java’s “garbage collector” which deletes objects that are no longer being used. To sidestep these memory issues, most of the objects generated internally by HAL are recycled rather than discarded. This reuse also has a performance benefit: if a function using the same object is called many times sequentially, the object will be faster to access in the computer’s memory because it was already cached from the earlier calls.

A key example of this reuse: when agents die and are removed from the model during a simulation run, the removed agents are kept internally and will be returned again for re-initialization when a new agent is requested. Agent recycling ensures that the number of agents that the model creates over a complete model run is capped to the maximum population that exists in the model at one time.

### 3.2 Component Overview

We now move from the abstract discussion of the unifying principles behind HAL to a look at its core components in more detail. Although it may seem that learning how to use these components would be a difficult task given their number and variety, all components were designed with a consistent API (Application Programming Interface), which makes changing between agent/grid types and learning their methods much easier.

#### 3.2.1 AgentGrids

AgentGrids are used as spatial containers for agents. They come in 1D, 2D, 3D, and non-spatial types. An example usage of a 3D AgentGrid is shown in Fig 3. These objects hold populations of agents that exist either bound to a lattice, or are free to move continuously. Internally, AgentGrids are composed of two data-structures: an agent list for agent iteration, and an agent lattice for spatial queries (even off-lattice agents are stored on a lattice for quick access). The agent list can be shuffled at every iteration to randomize iteration order, and the list holds onto removed agents to facilitate object recycling.

**Figure 3.**
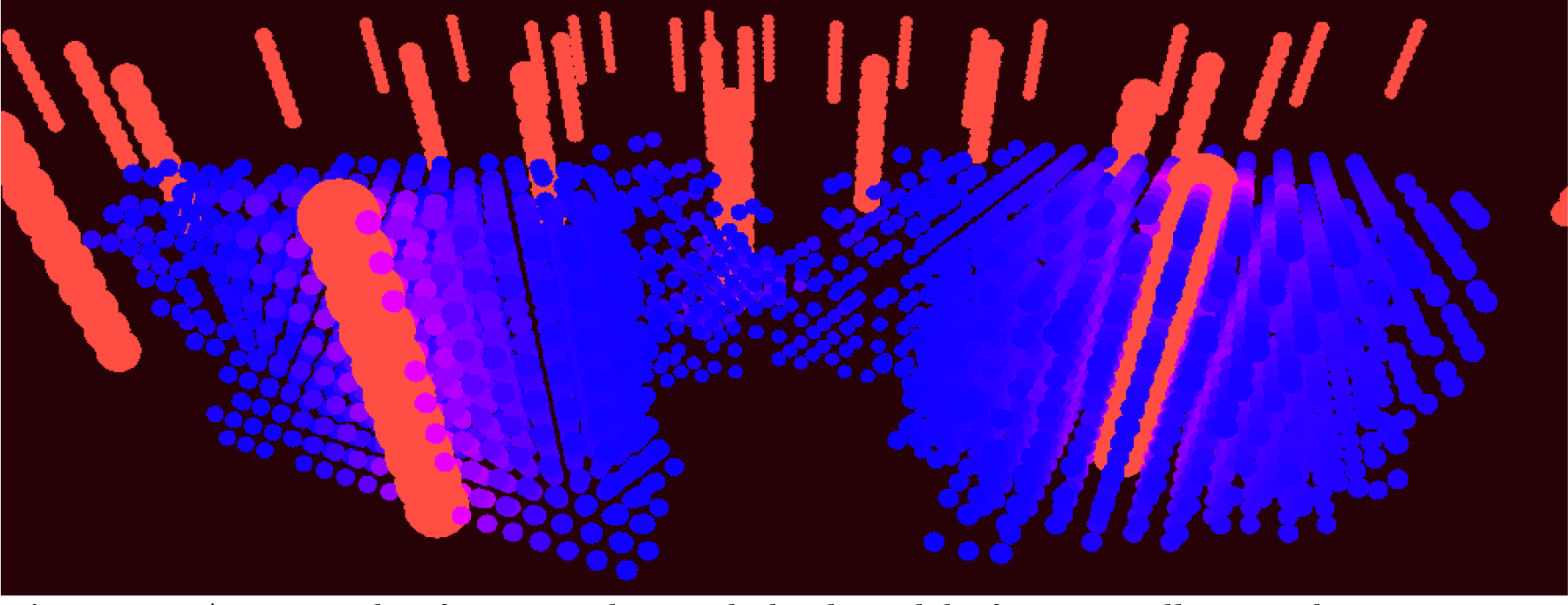
An example of a 3D on-lattice hybrid model of tumor cells spreading through tissue. The red vertical lines represent vessels, and the blue dots represent tumor cells. The cell color goes from pink to blue depending on how much oxygen is locally available. Displayed using the OpenGL3DWindow object.

#### 3.2.2 Agents

There are 10 base types of agent, introduced in Table 2. The SQ and PT suffixes refer to whether the agents are imagined to exist as lattice bound squares/voxels, or as non-volumetric points in space.

**Table 2.**
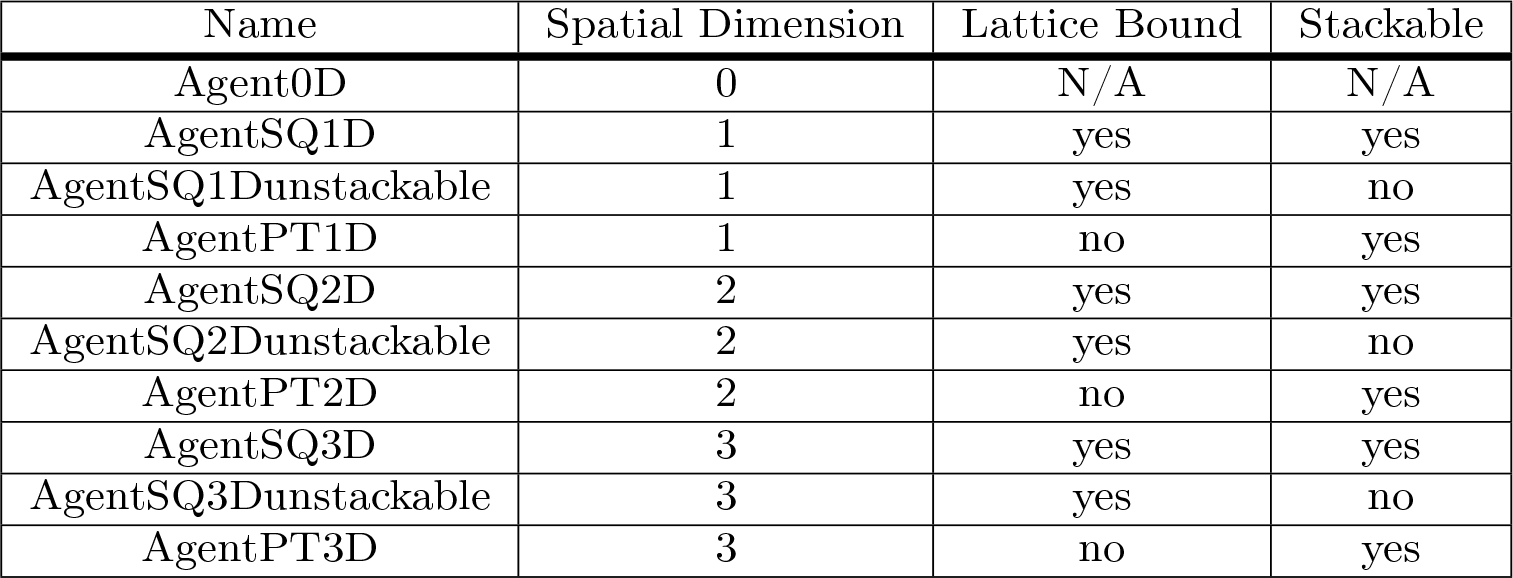
The 10 base agent types in HAL. The differences between them are displayed in each column. Stackable refers to whether multiple agents can exist on the same lattice position

Agent objects are always bound to a grid. In their base class form, agents keep track of their position on the grid and their age. Newly created agents are not included in the same iteration loop in which they are created, to prevent infinite loops of “runaway proliferation.” The base agent classes can be extended to include additional state properties and methods as needed.

#### 3.2.3 PDEGrids

The PDE Grids consist of either a 1D, 2D, or 3D lattice of concentrations. PDE grids contain functions that will solve reaction-diffusion equations. PDE function operations are accumulated on a separate lattice so they can be applied all at once in a simultaneous update. Currently implemented PDE solution methods include:

- Forward difference in time and 2nd order central difference in space diffusion
- ADI Diffusion [25]
- 1st order upwind finite difference advection for incompressible flows [26]
- 1st order finite volume upwind advection for compressible flows
- Modification of values at single lattice positions to facilitate reaction with agents or other sources/sinks.

Most of these methods are flexible, allowing for spatially heterogeneous diffusion rates and advection velocities as well as different boundary conditions such as periodic, Dirichlet, and zero-flux Neumann.

#### 3.2.4 Graphical User Interface (GUI)

The GUI building system consists of the following components:

- UIWindow: a window that displays GUI sub-components which are added in columns. the UIWindow will automatically scale to the appropriate size to fit all sub-components. The following five sub-components can be added:
  – UIGrid: a grid of pixels whose values are set individually. These are typically used to plot agent positions and diffusible concentrations, and can be easily output in GIF or PNG formats.
  – UIPlot: an extension of the UIGrid, the UIPlot is used to create real-time plots. The UIPlot will automatically resize to fit points that fall out of its bounds.
  – UILabel: a label that presents modifiable text.
  – UIButton: a button that executes a function when clicked
  – UIInputFields: various fields that accept bounded input of Integers, Doubles, Strings, Booleans, File Creation/Selection, and Combo boxes
- Window2DOpenGL/Window3DOpenGL: visualization windows that use OpenGL to efficiently render polygon graphics.
- GridWindow: A shortcut to generate a UIWindow with a single UIGrid component embedded. This simple component is used in the results section example.
- GifMaker: An object that can turn UIGrid visualization snapshots into gifs (Original source code created by Patrick Meister [27]).

An example GUI that uses the UIWindow with embedded UIButtons, InputFields, UILabels, and a UIGrid is shown in Fig 4.

**Figure 4.**
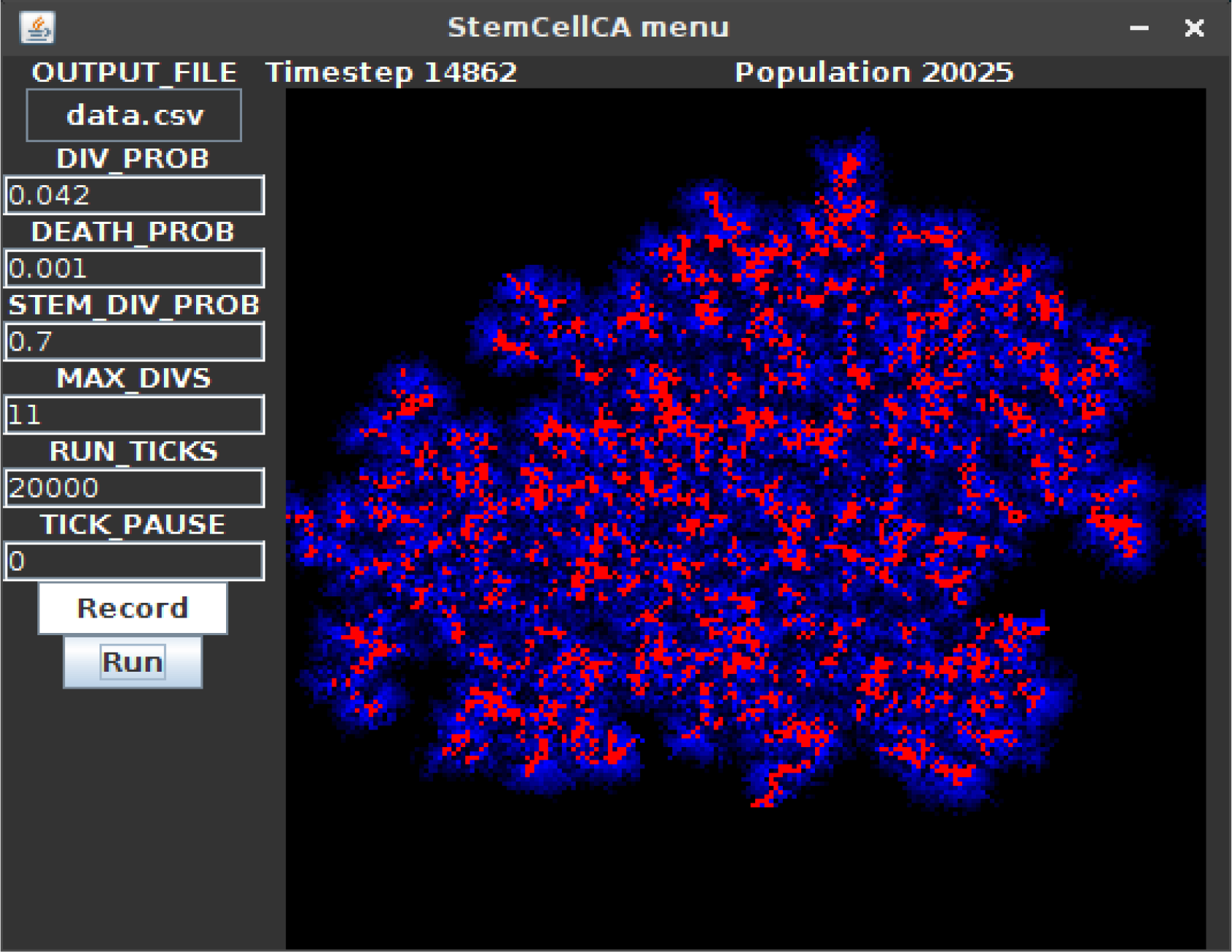
An example UIWindow GUI. When the “Run” UIButton (bottom left) is clicked, the UIGrid component (right) displays a running model that is parameterized with the given UIInputField settings (left). In this example model based on [28], the red cells are stem cells, and the blue cells are differentiated cells. Differentiated cells have a limited number of divisions and therefore can only spread a limited distance from the stem cells. UILabels (top) show the current timestep and population size.

##### 3.2.5 Utilities

The Util and Rand classes are used with almost every project. The Util class consists of a collection of standalone functions that solve common problems such as: Generating colors for use with the visualization tools, array manipulation, generating coordinate neighborhoods (eg. VonNeumann, Moore, Hex, Triangular), spatial mathematical operations, multicore parallelization, functions to save and load model states, etc. The Rand class is used for generating random numbers and for sampling distributions (eg. Gaussian, Poisson, Binomial, Multinomial - created using code adapted from the Colt and Numpy open source libraries [29, 30]) See the manual for more information [24].

##### 3.2.6 Tools

A set of miscellaneous tool objects are included to help with specific modeling tasks, these include:

- A FileIO object that is used to read input files and output results.
- A GenomeTracker object that internally stores phylogeny information in a searchable tree structure, and can be used to model branching processes.
- An ODESolver object that can solve ODEs numerically using Euler, Runge-Kutta 4, and Runge-Kutta Fehlberg 4,5 integration.
- A Multi-Well Experiment object that uses multi-threading to run and display many models simultaneously. The modeler simply creates an array of initialized models, defines an update and draw step, and can then feed many models into the Multi-Well experiment object and observe divergences in dynamics. This allows modelers to intuitively seed different models or replicates of the same conditions and observe differences in their behavior over time, see Fig 5
- An InteractiveModel object that embeds models in a graphical user interface from which the modeler can schedule modifications to parameters, such as treatment application, and interact with their model in real time. Modelers may also rewind execution to adjust settings, helping them to more quickly understand their model dynamics, and identify useful drug combinations and schedules. This tool exemplifies the power of modular design, and uses a UIPlot object for the timeline, as well as several UIGrids and UIButtons for other interactive components. This tool was used as part of the development of the Cancer Crusade game [31] to test the effects of therapy on a model by Tessi et al. [12], see Fig 6.

**Figure 5.**
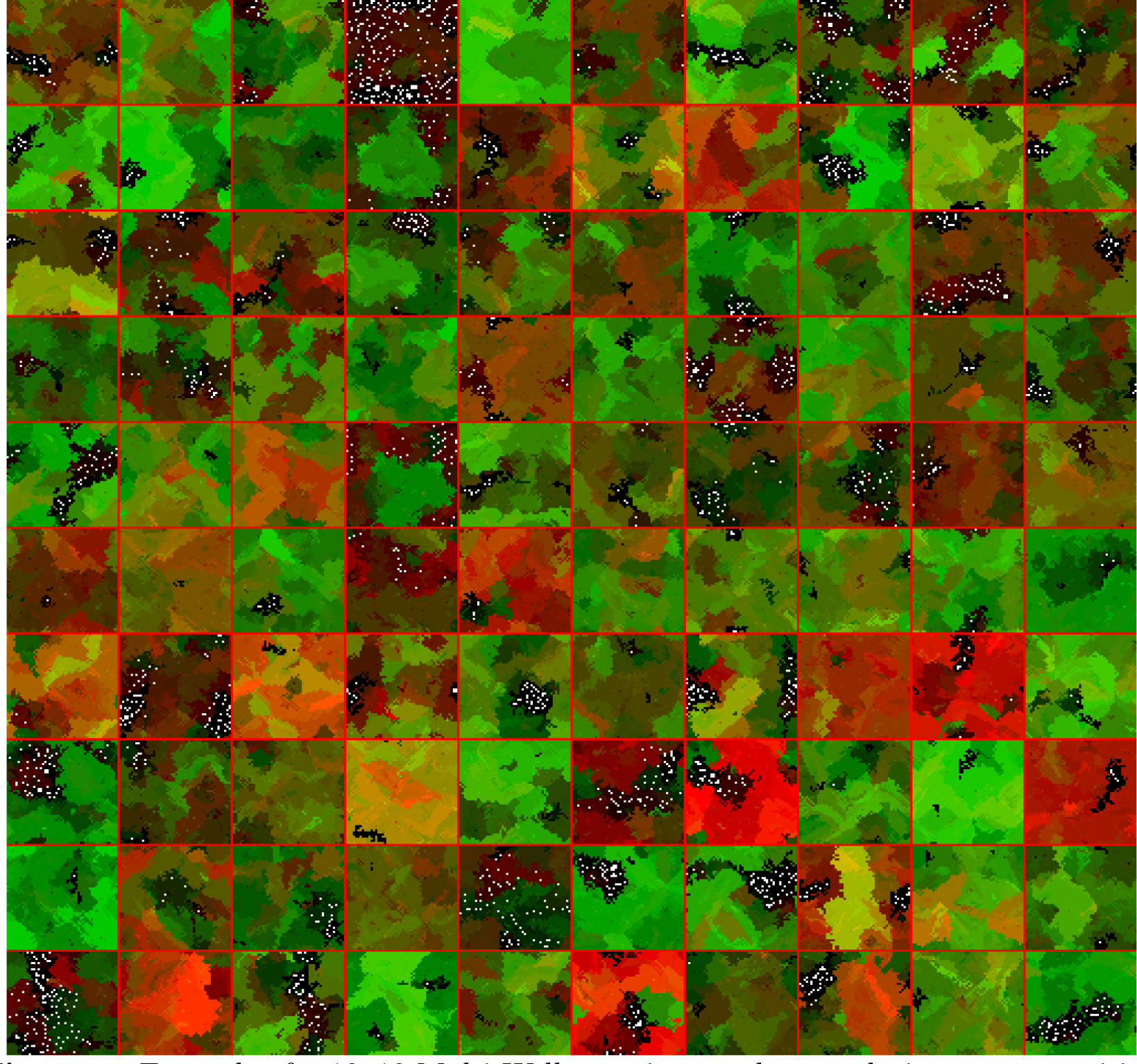
Example of a 10×10 Multi-Well experiment where evolutionary competition of two phenotypes (red,green) shows divergent results with different random seeds. models are separated with red lines.

**Figure 6.**
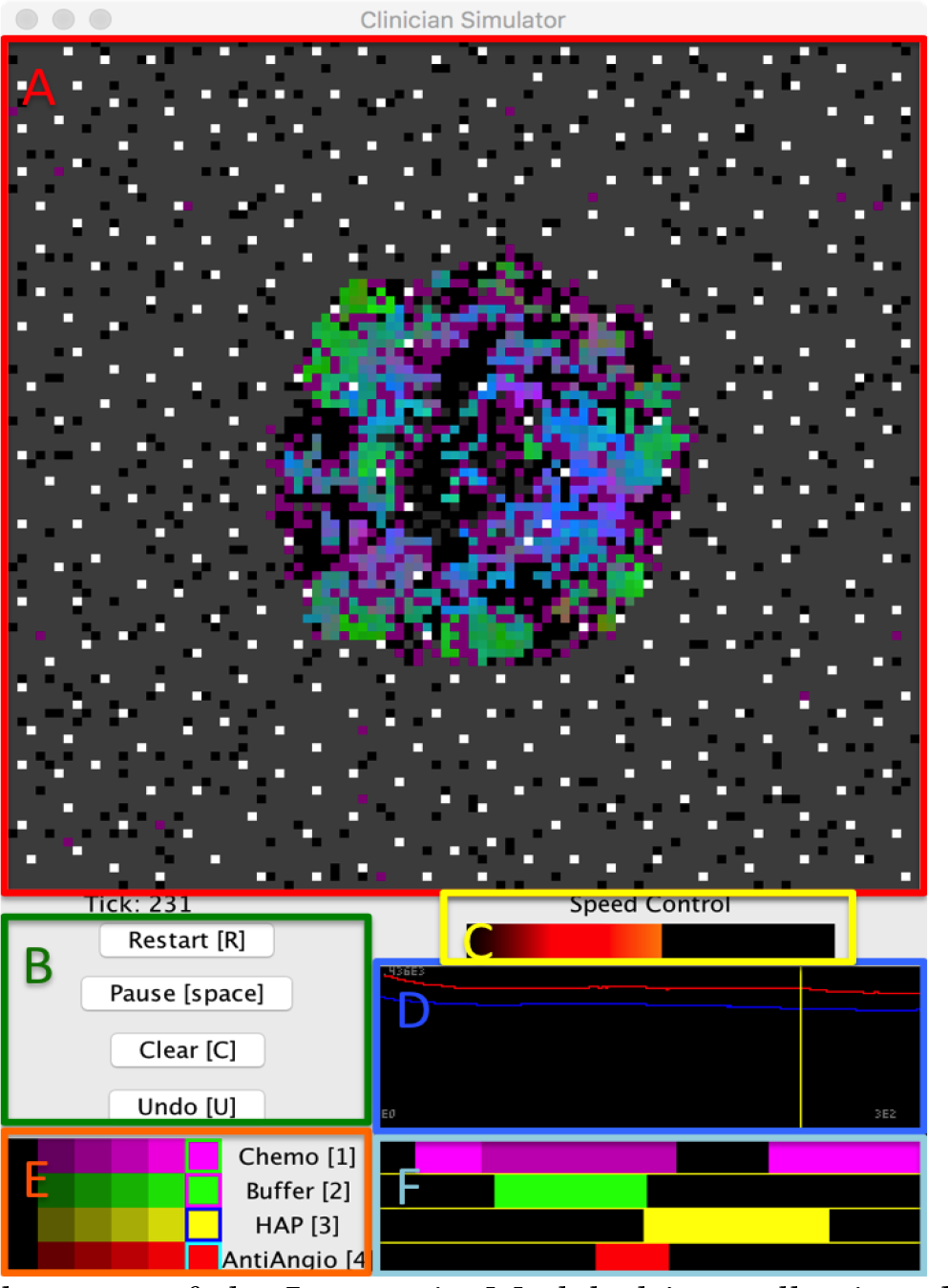
Example usage of the InteractiveModel object, allowing the modeler to experiment with treatment strategies using a model of the by Tessi et al. [12]. **(A)** A spatial visualization of the current model state. **(B)** A control panel of UIButtons allows the user to quickly restart the model, pause execution, clear all treatments, and undo previous changes. Hotkeys for these controls are in brackets. **(C)** A Speed Control option allows the user to easily adjust the execution speed of the model to range from evaluating as fast as possible to taking a second between time-steps, allowing for careful observation of model dynamics. **(D)** A timeline that will plot timestep information so that the user may observe aggregate changes over time in response to treatment. The user may also click anywhere in the timeline to backtrack to a previous timepoint and replay the model from there. The timeline will also automatically backtrack to recalculate any necessary frames when a treatment schedule change is made. **(E)** A set of sliders that allow the user to select different treatment intensities for each drug. **(F)** Each horizontal bar parallels the simulation timeline and displays the schedule of a different treatment. Modelers can click on regions within these bars to change regions to a new treatment intensity selected in (E). Modelers may also use the hotkeys presented in (E) to apply the selected intensities in real time as the model runs.

## 4 Results: Competitive Release Model

To demonstrate how the aforementioned principles and components of HAL are applied, we consider a simple but complete example of hybrid modeling. We implement the model of pulsed therapy based on a recent publication by Gallaher et al. [14]. We also showcase the flexibility that the modular component approach brings by displaying three different parameterizations of the same model side by side.

### 4.1 Competitive Release Introduction

The model in [14] describes two competing tumor-cell phenotypes: a rapidly dividing, drug-sensitive phenotype and a slower dividing, drug-resistant phenotype. There is also a diffusible drug that enters the system through the domain boundaries and is consumed by the tumor cells over time.

Every timestep (“tick”) each cell has a probability of death and a probability of division. The division probability depends on phenotype (resistant cells divide less frequently) and the availability of space (cells will divide only if there is an open space in the nearest eight grid square neighborhood or moore neighborhood). Sensitive cells have a death rate that increases when the cells are exposed to drug, while resistant cells have a constant death rate.

The modular design of HAL allows us to test 3 different treatment conditions, each with an identical starting tumor (no drug, constant drug, and pulsed drug). An interesting outcome of the experiment is that pulsed therapy is better at managing the tumor than constant therapy. Under pulsed therapy the sensitive population is kept in check, while still competing spatially with the resistant phenotype and preventing its expansion. The rest of the section describes in detail how this abstract model is generated.

Fig 7 provides a high level look at the structure of the code. Red font indicates where a section of the coding example is called. Table 3 provides a quick reference for the built-in HAL functions used in this example. Any functions that are used by the example but do not exist in the table are defined within the example itself and explained in detail below the code. Those fluent in Java may be able to understand the example just by reading the code and using Table 3. Built-in HAL functions and classes are highlighted in red in the following source code to make identifying HAL’s components easier.

**Figure 7.**
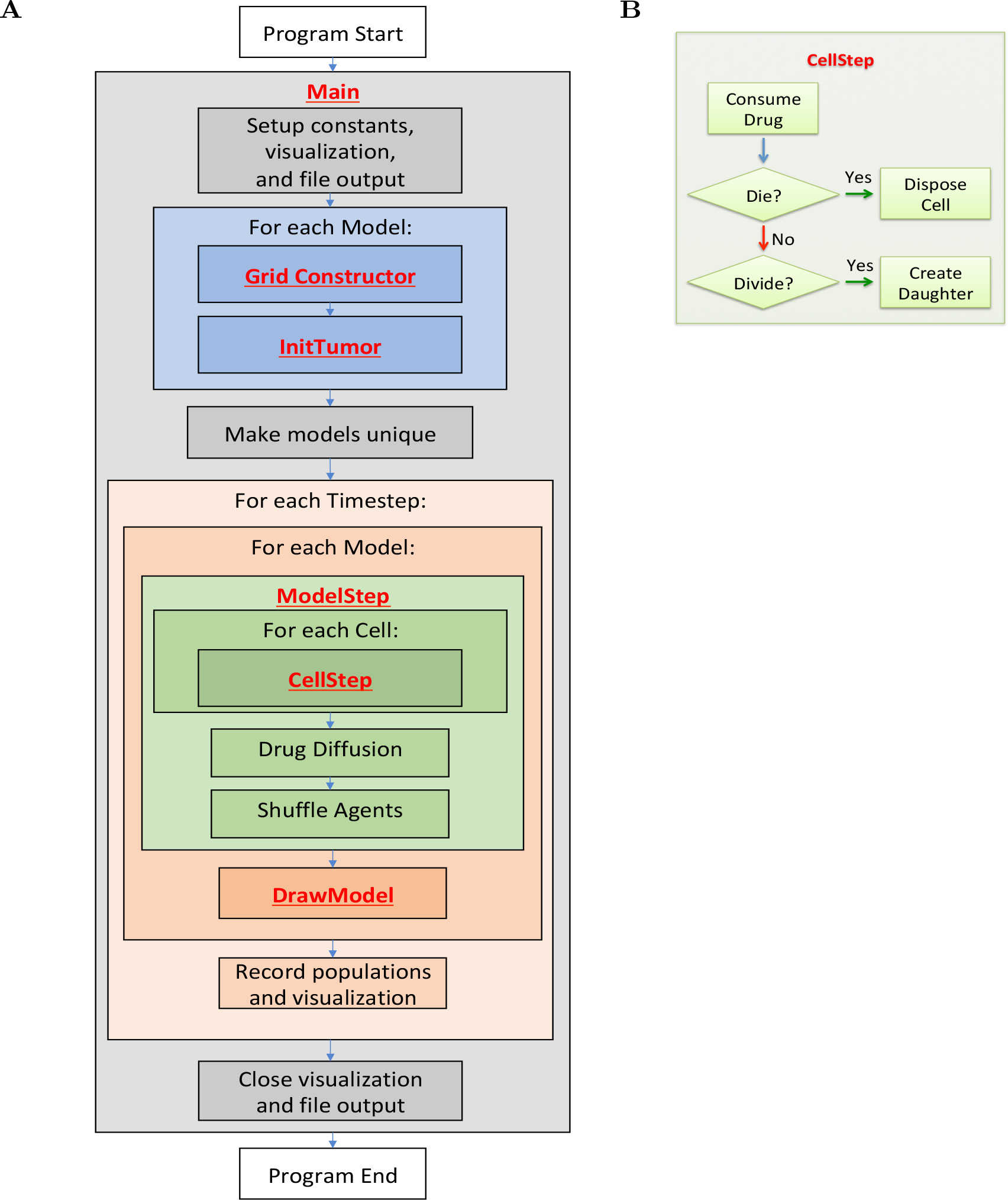
**(A)** Example program flow diagram. Red font indicates where coding example sections are first called. **(B)** CellStep function flow diagram.

**Table 3.**
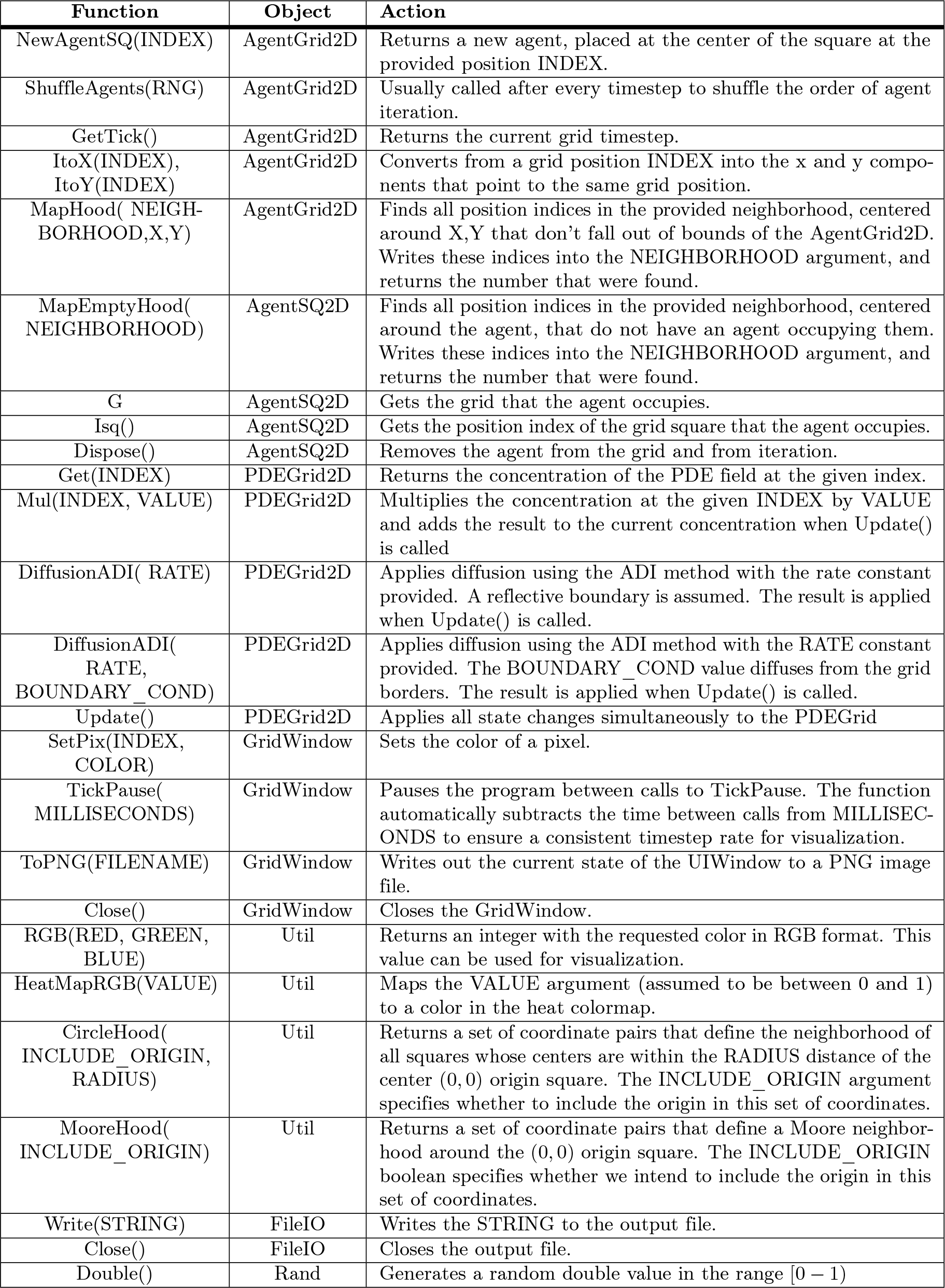
HAL functions used in the example. Each function is a method of a particular object, meaning that when the function is called it may use properties that pertain to the object that it is called from.

### 4.2 Main Function

We first examine the ‘main’ function for a bird’s-eye view of how the program is structured. Source code elements highlighted in red are built-in HAL functions and objects, and can be referenced in Table 3.

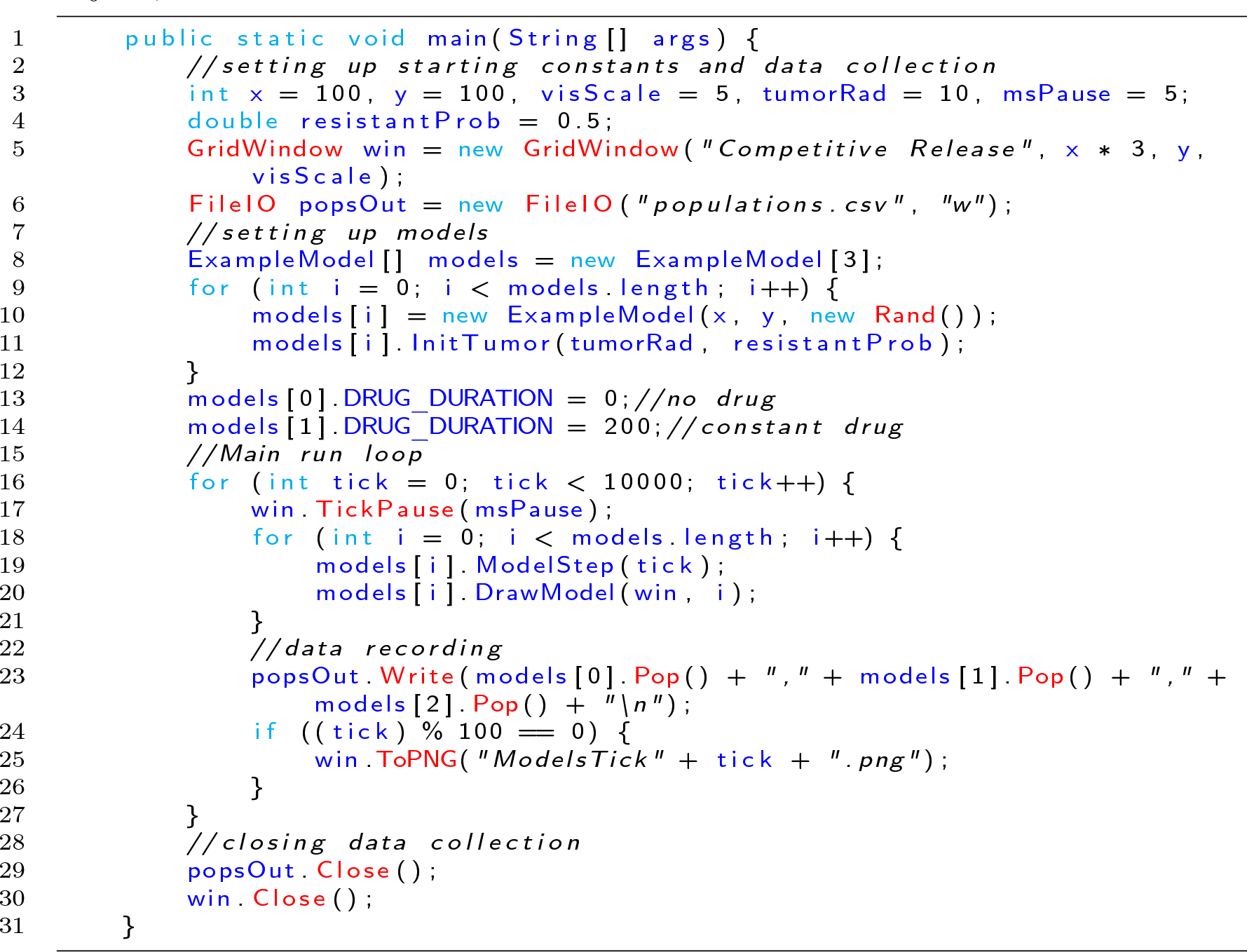

Lines 3-4: Defines all of the constants that will be needed to setup the model and display.
5: Creates a GridWindow of RGB pixels for visualization and for generating timestep PNG images. *3, y define the dimensions of the pixel grid. the x variable is multiplied by 3 so that 3 models can be visualized side by side in the same window. The last argument is a scaling factor that specifies that each pixel on the grid will be viewed as a 5×5 square of pixels on the screen.
6: Creates a file output object that will write to a file called populations.csv.
8: Creates an array with 3 entries that will be populated with models.
9-12: Fills the model list with models that are initialized identically. Each model will hold and update its own cells and diffusible drug. See the Grid Definition and Constructor section and the InitTumor Function section for more details.
13-14: Setting the DRUG_DURATION constant creates the only difference in the 3 models being compared. In models[0] no drug will be administered. In models[1] drug administration will be constant (DRUG_DURATION is set equal to DRUG_CYCLE). In models[2] drug will be administered periodically (the default value of DRUG_DURATION is 40). See the ExampleModel Constructor and Properties section for the default model initialization.
16: Executes the main loop for 10000 timesteps.
17: Requires every iteration of the loop to take a minimum number of milliseconds. This slows down the execution and display of the model and makes it easier for the viewer to follow.
18: Loops over all models to update them.
19: Advances the state of the agents and diffusibles in each model by one timestep. See the Model Step Function for more details.
20: Draws the current state of each model to the window. See the Draw Model Function for more details.
23: Writes the population sizes of each model every timestep to allow the models to be compared.
24: Every 100 timesteps, writes the state of the model as captured by the GridWindow to a PNG file.
29-30: After the main for loop has finished, the FileIO object and the visualization window are closed, and the program ends.

### 4.3 ExampleModel Constructor and Properties

This section explains how the grid is defined and instantiated.

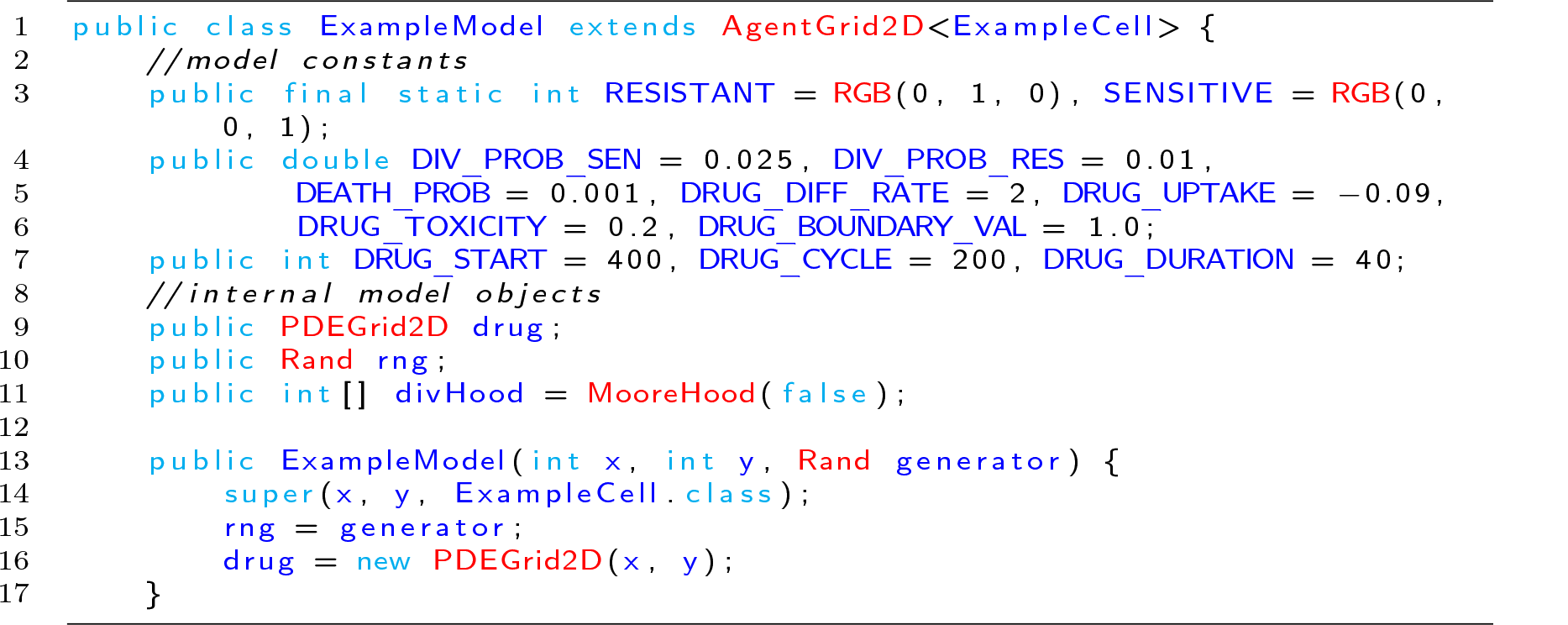

1: The ExampleModel class, which is user defined and specific to this example, is built by extending the generic AgentGrid2D class. The extended grid class requires an agent type parameter, which specifies the type of agent that will live on the grid. To meet this requirement, the <ExampleCell> type parameter is added to the declaration.
3: Defines RESISTANT and SENSITIVE constants, which are created by the Util RGB function. These constants serve as both colors for drawing and as labels for the different cell types.
4-7: Defines all constants that will be needed during the model run. These values can be reassigned after model creation to test different parameter settings. In the main function, the DRUG_DURATION variable is modified for the Constant-Drug, and Pulsed Therapy experiment cases.
9: Declares that the model will contain a PDEGrid2D, which will hold the drug concentrations. The PDEGrid2D can only be initialized when the x and y dimensions of the model are known, which is why we do not assign their values until the constructor function is called.
10: Declares that the Grid will contain a Random number generator (the Rand object), but takes it in as a constructor argument to allow the modeler to seed the generator if desired for consistent output.
11: Creates a neighborhood using the MooreHood function. The MooreHood function generates a set of coordinates that define the Moore Neighborhood (the 8 closest coordinates to a central origin), centered around (0, 0). The false argument declares that we do not want to include the origin in the neighborhood, just the 8 coordinates around that position. The neighborhood is stored in the format [0_1_0_2_,*…,* 0_*n*_, *x*_1_, *y*_1_, *x*_2_, *y*_2_,*…, x*_*n*_, *y*_*n*_]. The leading zeros are written to when MapHood is called, and will store the position indices that the neighborhood maps to. See the CellStep function for more information, and the InitTumor Function Line 3 for another example of the use of neighborhoods
13: Defines the model constructor, which takes as arguments the x and y dimensions of the model and a random number generator (a Rand object).
14: Calls the AgentGrid2D constructor with super, passing it the x and y dimensions of the model, and the ExampleCell Class. This Class is used by the Grid to generate a new cell when the NewAgentSQ function is called.
15-16: The Rand argument is assigned and the drug PDEGrid2D is defined with matching dimensions.

### 4.4 InitTumor Function

The next segment of code is a function from the ExampleModel class that defines how the tumor is first seeded after the ExampleModel is created.

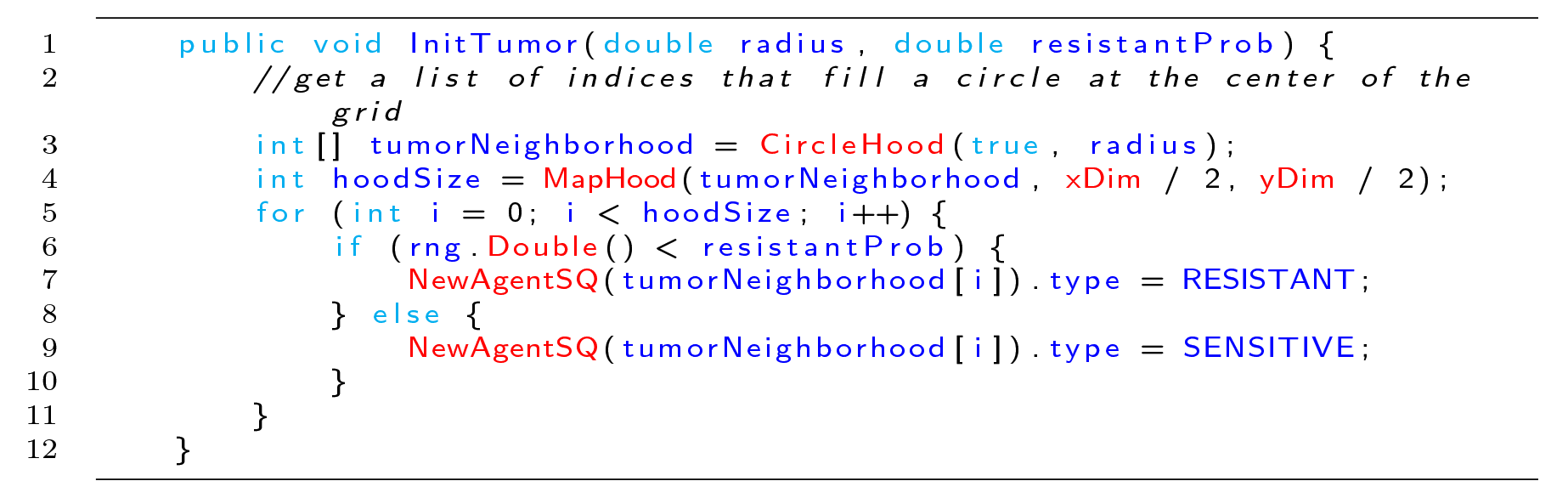

1: The arguments passed to the InitTumor function are the approximate radius of the circular tumor being created and the probability that each created cell will be of the resistant phenotype.
3: Sets the tumorNeighborhood array using the CircleHood function, which stores coordinates in the form [0_1_, 0_2_,*…,* 0_*n*_, *x*_1_, *y*_1_, *x*_2_, *y*_2_,*…x*_*n*_, *y*_*n*_]. The x,y coordinate pairs define a neighborhood of all squares whose centers are within the radius distance of the center (0, 0) origin square. The leading 0s are used by the MapHood function to store the mapped indices. The boolean argument specifies that the origin will be included in this set of squares, thus making a completely filled circle of squares.
4: Uses the MapHood function to map the neighborhood defined above to be centered around xDim/2,yDim/2 (the dimensions of the AgentGrid). The results of the mapping are written as position indices to the beginning of the tumorNeighborhood array. MapHood returns the number of valid indices found, and this will be the size of the starting population.
5: Loops from 0 to hoodSize, allowing access to each mapped position index in the tumorNeighborhood.
6: Samples a random number in the range [0 – 1) and compares to the resistantProb argument to set whether the cell should have the resistant phenotype or the sensitive phenotype.
7-9: Uses the NewAgentSQ function to place a new cell at each tumorNeighborhood position. In the same line we also specify that the phenotype should be either resistant or sensitive, depending on the result of the rng.Double() call.

### 4.5 ModelStep Function

This section looks at the model’s step function which is executed once per timestep by each Model.

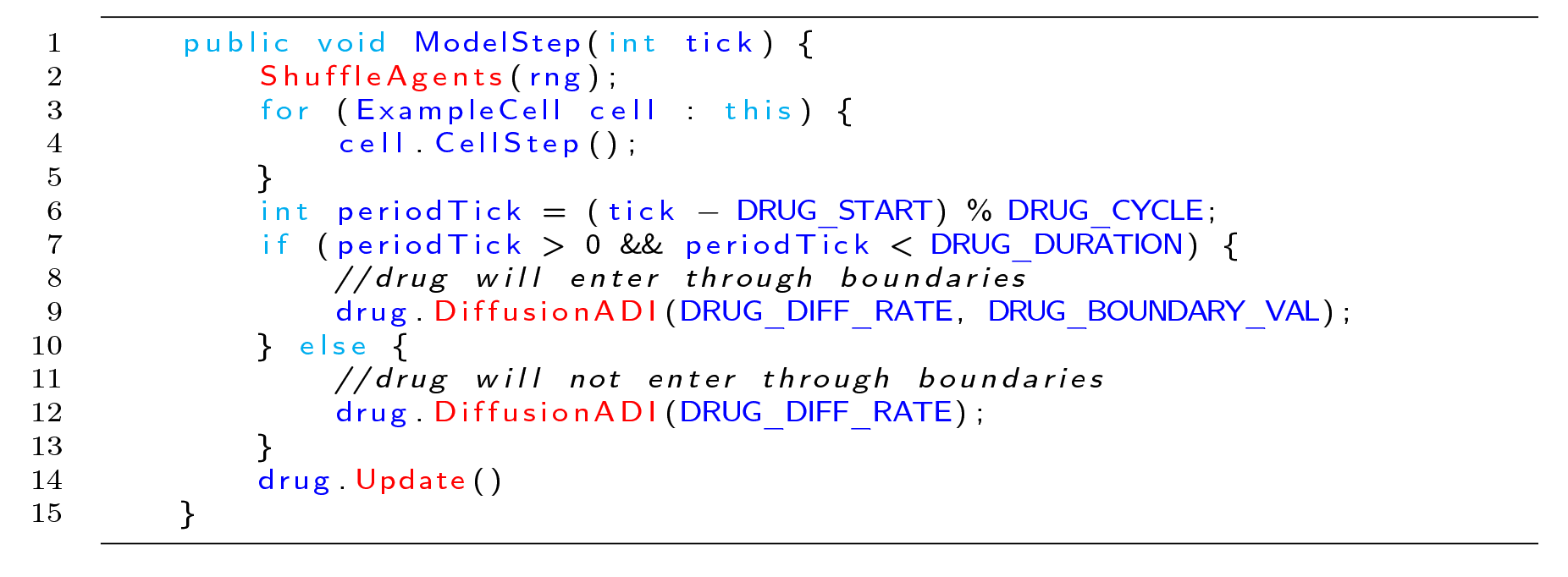

2: The ShuffleAgents function randomizes the order of iteration so that the agents are always looped through in random order.
3-4: Iterates over every cell on the grid, and calls the CellStep function on every cell.
6-7: The periodTick variable stores at what point in the drug delivery cycle the tick is, and the If statement checks whether the tick is in the right part of the drug cycle to apply drug, (See the Grid Definition and Constructor section for the values of the constants involved, the DRUG_DURATION variable is set differently for each model in the Main Function)
9: If it is time to add drug to the model, the DiffusionADI function is called. DiffusionADI uses the ADI method which is more stable than 2D Euler and allows us to take larger steps. The additional argument to the DiffusionADI function specifies the boundary condition value DRUG_BOUNDARY_VAL. This causes the drug to diffuse into the PDEGrid2D from the boundary. Here we assume that drug is only delivered from the boundaries of the domain
12: Without the second argument the DiffusionADI function assumes a zero-flux boundary, meaning that drug concentration cannot escape or enter through the sides of the model. Therefore the only way for the drug concentration to decrease is via uptake by the Cells. See the CellStep function section, line 6, for more information.
14: Update is called to apply the reaction and diffusion changes to the PDEGrid.

### 4.6 CellStep Function and Cell Properties

We next look at how the ExampleCell Agent is defined and at the CellStep function that runs once per Cell per timestep. The G property that is referenced many times in this section is a built-in agent property that gives access to the ExampleGrid object that the cell lives on.

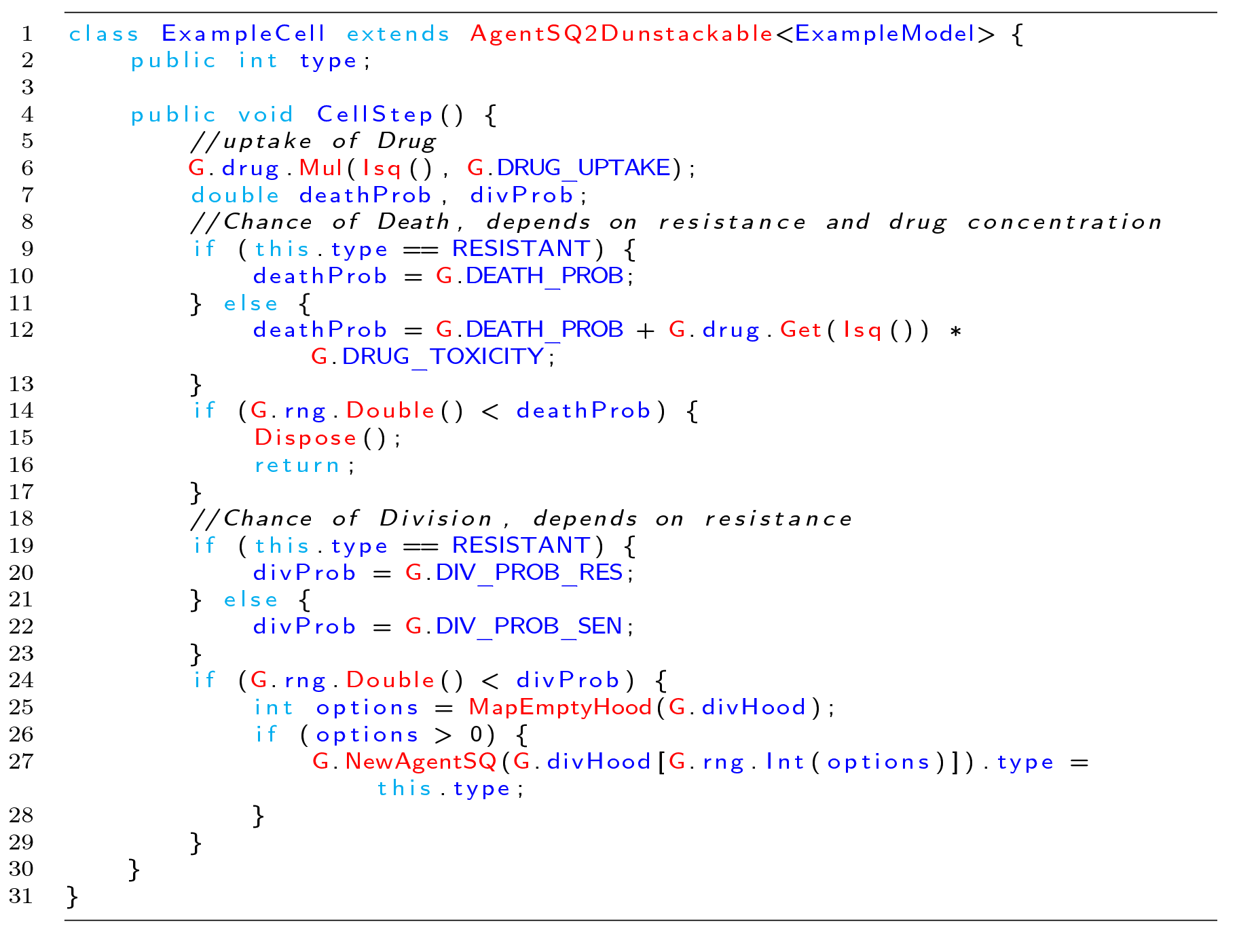

1: The ExampleCell class is built by extending the generic AgentSQ2Dunstackable class. The extended Agent class requires the ExampleModel class as a type argument, which is the type of Grid that the Agent will live on. To meet this requirement, we add the <ExampleModel> type parameter to the extension.
2: Defines a cell property called “type”. Each Cell holds a value for this field. If the value is RESISTANT, the Cell is of the resistant phenotype, if the value is SENSITIVE, the cell is of the sensitive phenotype. The RESISTANT and SENSITIVE values are defined in the ExampleGrid as constants (See the ExampleModel Constructor and Properties, line 3).
6: The G property is used to access the ExampleGrid object that the Cell lives on. G is used often with agent functions as the AgentGrid is expected to contain any information that is not local to the individual agent. Here it is used to get the drug PDEGrid2D. The drug concentration at the index that the Cell is currently occupying (Isq()) is then multiplied by the drug uptake constant, thus modeling local drug uptake by the Cell.
7: Defines deathProb and divProb variables, these will be assigned different values depending on whether the ExampleCell is RESISTANT or SENSITIVE.
9-12: If the cell is resistant, the deathProb variable is set to the DEATH_PROB value alone, if the cell is sensitive, an additional term is added to account for the probability of the cell dying from drug exposure, using the concentration of drug at the cell’s position (Isq())
14-16: Samples a random number in the range [0 – 1) and compares to deathProb to determine whether the cell will die. If so, the built-in agent Dispose() function is called, which removes the agent from the grid, and then return is called so that the dead cell will not divide.
19-22: Sets the divProb variable to either DIV_PROB_RES for resistant cells, or DIV_PROB_SEN for sensitive cells.
24: Samples a random number in the range [0 – 1) and compares to divProb to determine whether the cell will divide.
25: If the cell divides, the MapEmptyHood function is used, which checks the positions in the divHood (the Moore neighborhood as defined in the ExampleModel Constructor and Properties section, line 11) around the Cell, and writes the position indices that do not contain any agents into the divHood. MapEmptyHood returns the number of valid empty positions found.
26-27: If there are one or more valid division options, a new daughter cell is created using the NewAgentSQ function and its starting location is chosen by randomly sampling the divHood array to pull out one if its valid locations. The other daughter is assumed to occupy the same location as the mother cell. Finally with the.type=this.type statement, the phenotype of the newly placed daughter cell is inherited from the mother cell.

### 4.7 DrawModel Function

We next look at the DrawModel Function, which is used to display a summary of the model state on a GridWindow object. In this program, DrawModel is called once for each model per timestep; see the main function section for more information.

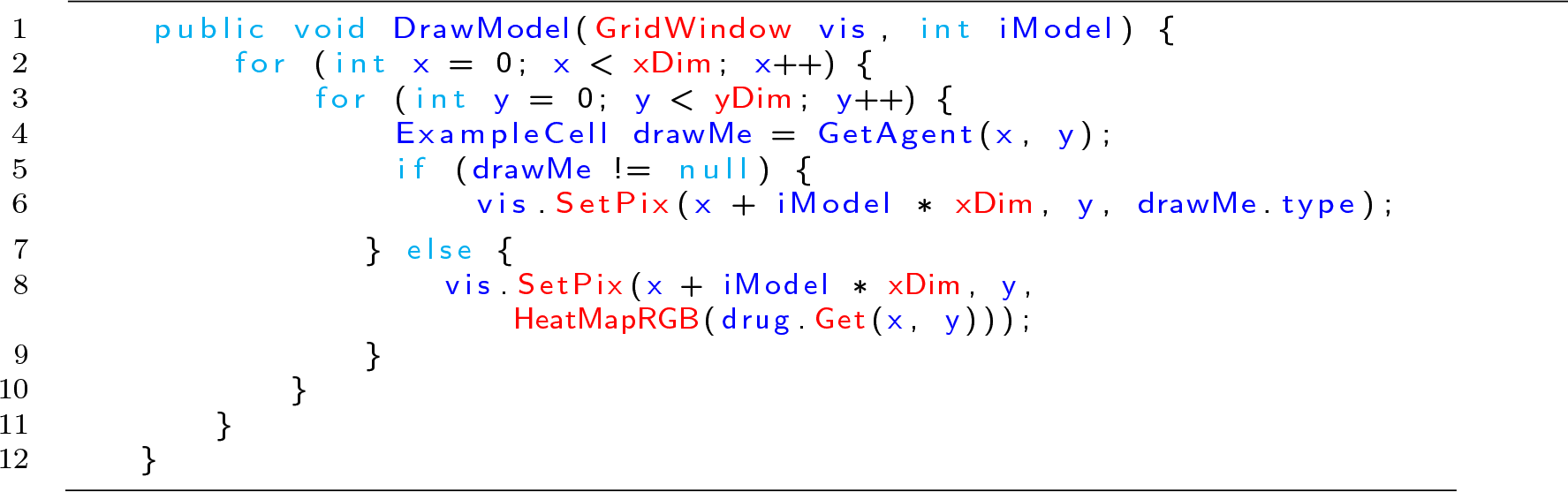

2-3: Loops over every lattice position of the grid being drawn, xDim and yDim refer to the dimensions of the model.
4: Uses the GetAgent function to get the Cell that is at the x,y position.
5-6: If a cell exists at the requested position, the corresponding pixel on the GridWindow is set to the cell’s phenotype color. To draw the models side by side, the pixel being drawn is displaced to the right by the model index.
7-8: If there is no cell to draw, then the pixel color is set based on the drug concentration at the same index, using the built-in heat colormap.

### 4.8 Imports

The final code snippet looks at the imports that are needed. Any modern Java IDE should generate import statements automatically.

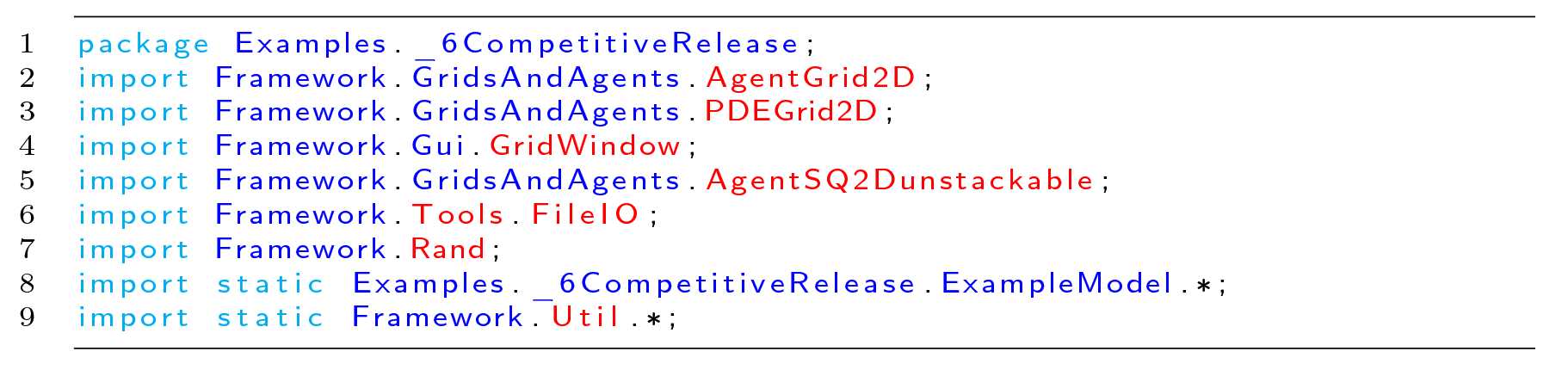

1: The package statement specifies where the file exists in the larger project structure
2-7: Imports all of the classes that we will need for the program.
8: Imports the static fields of the model so that we can use the type names defined there in the Agent class.
9: Imports the static functions of the Util file, which adds all of the Util functions to the current namespace, so we can natively call them. Statically importing Util is recommended for every project.

### 4.9 Model Results

Table 4 displays the model visualization at timestep 0, timestep 400, timestep 1100, timestep 5500, and timestep 10,000 recorded from the GridWindow ToPNG function. The caption explores the notable trends visible in each image. Fig 8 displays the population sizes as recorded by the FileIO Write function at the end of every timestep.

**Table 4.**
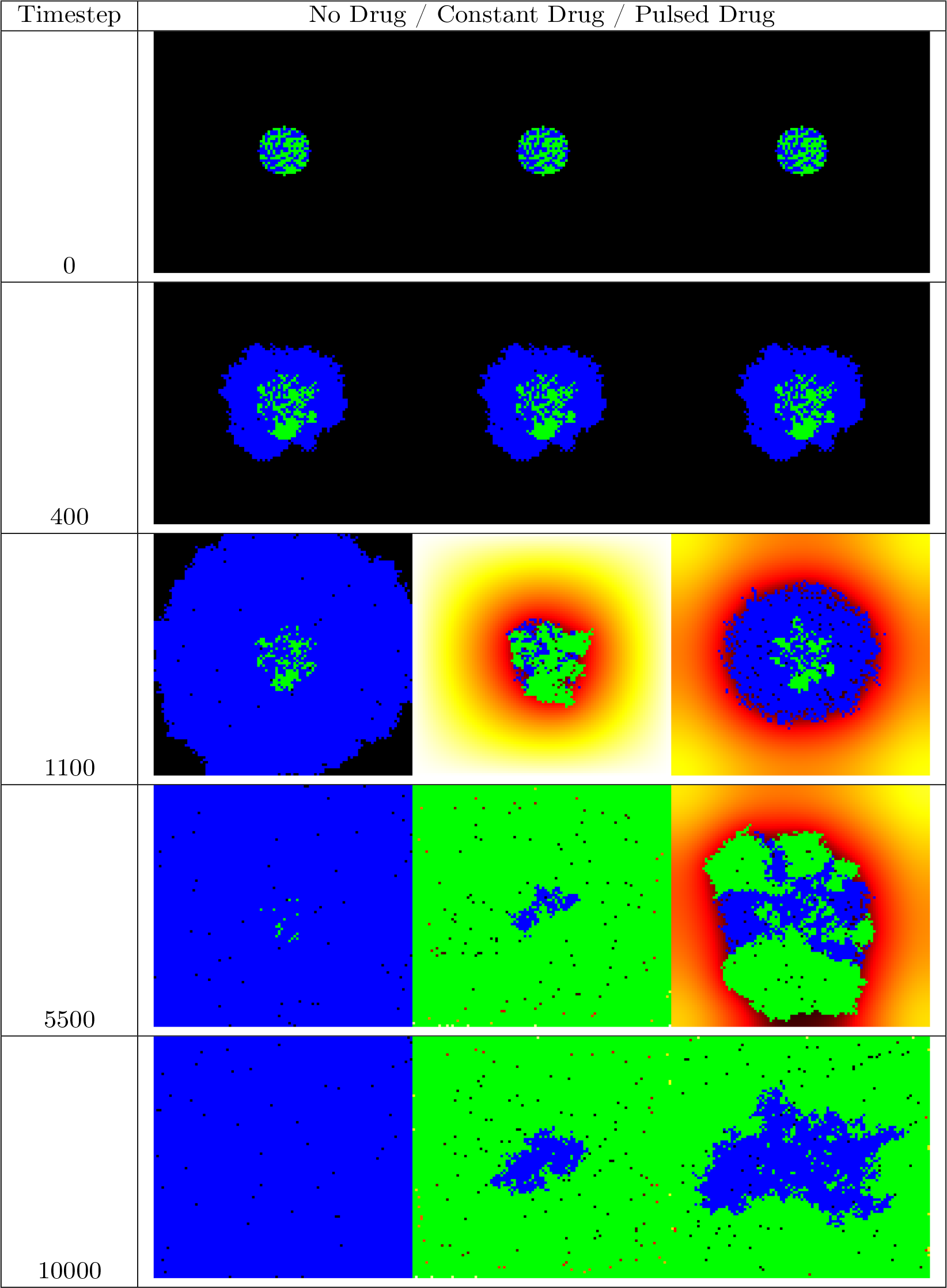
Selected model visualization PNGs. Blue cells are drug sensitive, Green cells are drug resistant, background heatmap colors show drug concentration.

**Figure 8.**
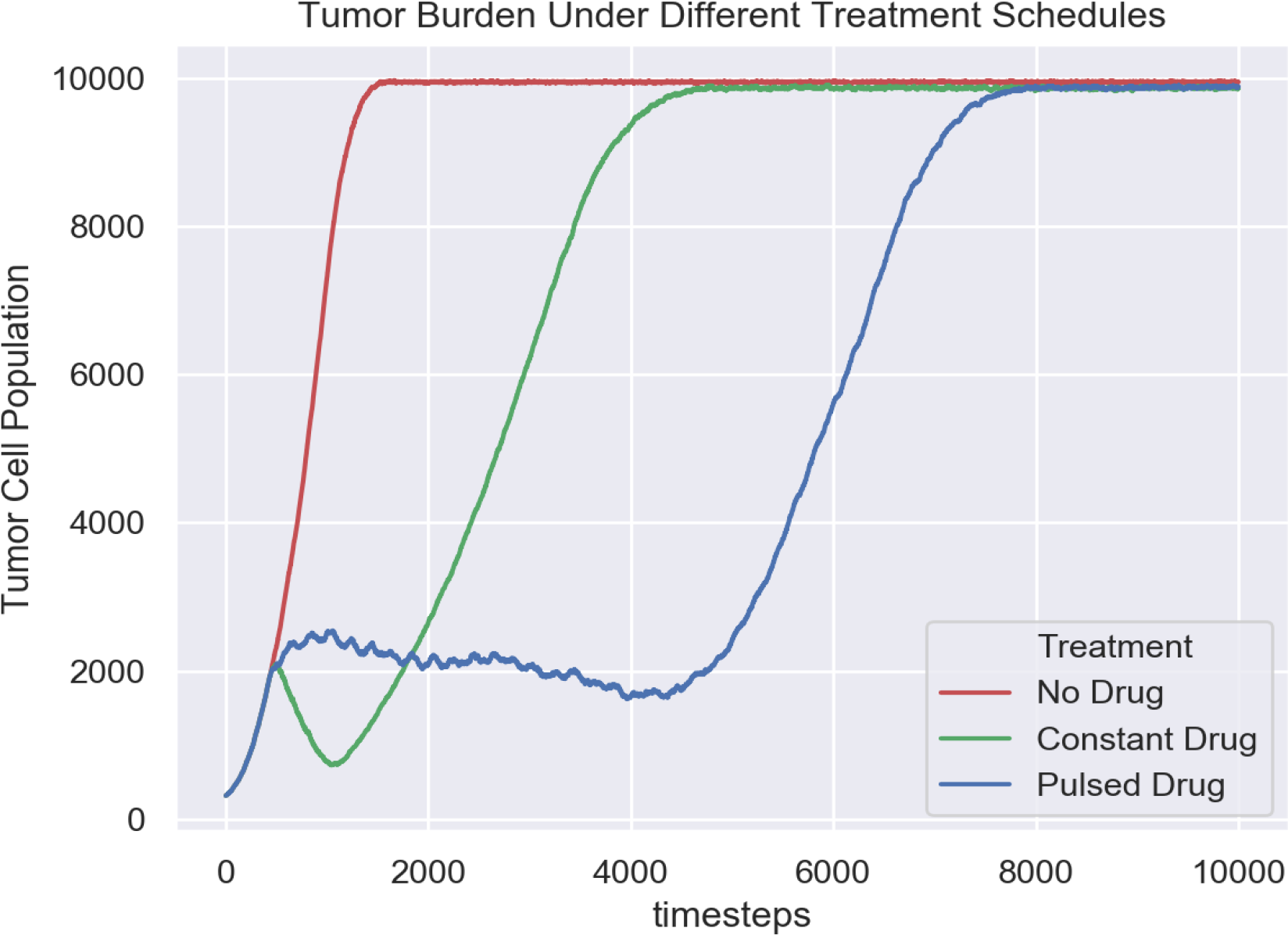
FileIO population output. This plot summarizes the changes in tumor burden over time for each model. This plot was constructed in python using data accumulated in the program output CSV file. Displayed using Seaborn with Python.

This example illustrates the power of HAL’s approach to model building. Writing relatively little complex code, we setup a three model experiment with nontrivial dynamics along with methods to collect data and visualize the models. We now briefly review the model results.

As can be seen in Table 4, at timestep 0 and timestep 400 (right before drug application starts), all 3 models are identical. At timestep 1100 the differences in treatment application show different effects: when no drug is applied, the rapidly dividing sensitive cells quickly fill the domain, while when drug is applied constantly, the resistant cells overtake the sensitive population. Pulsed drug kills some sensitive cells, but leaves enough alive to prevent growth of the resistant cells. At timestep 5500, the resistant cells have begun to emerge from the center of the pulsed drug model. At timestep 10000, all domains are filled. Interestingly, in the models with drug application, the sensitive cells are able to survive in the center of the domain because drug is consumed by cells on the outside. This creates a drug-free zone in which the sensitive cells out-compete the resistant cells even when drug is applied constantly.

As can be seen in Fig 8, the pulsed therapy is the most effective at preventing tumor growth, however the resistant cells ultimately succeed in breaking out of the tumor center and out-competing the sensitive cells on the fringes of the tumor. It may be possible to contain a population of sensitive and resistant cells for longer by using a different pulsing schedule or by modifying the treatment schedule in response to the tumor growth (adaptive therapy). As the presented model is primarily an example, we do not explore how to improve treatment further. For a more detailed exploration of the potential of adaptive therapy for prolonging competitive release, see [14].

## 5 Availability And Future Directions

### 5.1 How to Download and Contribute

HAL is publicly available on GitHub, at https://github.com/torococo/HAL. A manual is included that walks the user through installation and serves as a coding reference. For those interested in using HAL, downloading and setting up the project is a good first start. From there running and examining the included examples is recommended, as they do a good job of summarizing HAL’s capabilities. Modelers can contribute tools that they develop by making pull requests to the repository.

### 5.2 Future Directions

#### 5.2.1 Additional agent-based Modeling Paradigms

Currently the only paradigm implemented on top of the base agent types are the SphericalAgent2D/3D extension classes, which facilitate modeling cells as circles/spheres with Newtonian physics. In the future we hope to incorporate additional modeling paradigms that are commonly used in agent-based modeling of cells. A potential addition is a Delaunay Agent type, which would use Delaunay tessellation [32] to find the cell’s nearest neighbors and determine cell volume. We are also considering including modeling paradigms that construct cells out of smaller subunits, such as Deformable Ellipsoid Cell Modeling [33], as this would allow us to model the mechanics of tissue formation and cell migration in more detail.

#### 5.2.2 Cross Model Validation

Having many different paradigms to choose from adds several complications to modeling: It can take significant effort to build a model from scratch under one paradigm, and then significant additional effort to migrate the model to a different paradigm. By adding more modeling approaches with a consistent interface, HAL will lower the model migration barrier and allow modelers to test the merits of many paradigms in their investigation, and to validate their results by seeing whether they hold true across paradigms. Note that our goal is not to recreate all of the functionality of the pre-existing frameworks that support these paradigms, it is to provide their core algorithms so that users can easily choose from and compare them.

#### 5.2.3 Bridging Spatial Scales

We also hope to explore the possibility of adjusting spatial scales for both our PDEs and Agents. For PDEs, this is a readily understood problem, and we intend to add scalable PDEGrids to HAL soon. However, for agent-based modeling the process of changing scales while preserving dynamics is not so well defined, though we imagine that it may be possible under certain assumptions. This would be useful for helping us bridge the divide between cell level and tissue/organ/tumor level dynamics, as the number of cells involved at these scales are orders of magnitude greater than what desktop machines can tractably model.

#### 5.2.4 Assumption Modules

A common modeling task is exploring how combinations of different assumptions influence model behavior. The included ModuleSetManager object helps design models specifically with this in mind. The design entails providing code “hooks” so that code can be added to influence specific agent decisions and model events, (eg. whether an agent will reproduce). Modelers can then write assumption modules that will influence these events (eg. by altering the probability of reproduction based on an environmental factor that would otherwise be ignored).

This approach allows modelers to combine and remove assumption modules without having to worry about breaking the model. This facilitates easy exploration of the space of assumptions until ones suitable for understanding biological phenomena are found. We are very excited about the potential of this approach for collaborative projects and for building increasingly complex models by encapsulating the complexity into manageable parts, and hope to improve on the tools for this paradigm as we explore its potential.

#### 5.2.5 Advanced Scheduling

Taking inspiration from Repast, SWARM, and MASON, another expected extension is the inclusion of optional schedulers to facilitate more complex methods of iterating through agents than simply looping over each grid. This is not intended to replace the simple grid iteration approach, but instead should augment it with optional complex methods. An AgentList object is currently included to begin to address this. It allows modelers to make selective lists of agents for more flexible iteration.

#### 5.2.6 Building a Community

HAL has already seen adoption within the labs at the Integrated Mathematical Oncology department of Moffitt Cancer Center and beyond. We certainly hope that more outside users will be interested in its potential. As the user-base for HAL grows, we plan to extend the base of resources around the platform. The current set of resources that exist for new users to get started are the manual 6, a website with an online version of the manual [1] and a playlist of YouTube videos [34]. We intend to increase HAL’s online presence by including a website with a code repository to make sharing models and tools easier.

## 6 Conclusion

Cancer is a complex and heterogeneous disease whose mathematical study is still being developed. To make better progress in this endeavor, it is helpful to have a set of highly generic tools that encapsulate the key components of spatial modeling so that researchers can produce efficient models quickly without being constrained in their approach, nor in the complexity of the systems that they can produce. HAL is our attempt to achieve this.

HAL was made easily extensible so that researchers can build models and more specific tools on top of HAL’s generic base. The hope is that by this process HAL will grow into a powerful toolset that will help standardize and coordinate hybrid modeling in mathematical oncology.

We recommend HAL to anyone building spatial models for oncology, as the tools presented are primarily geared toward this end. However, given the amount of overlap and cross talk between the approaches used in different modeling applications, we hope that modelers outside of mathematical oncology will also take interest and contribute, to our mutual benefit.

## Supporting information

Supplemental Figure 1: HAL Manual

## Supporting information

**S1 Fig. HAL (Hybrid Automata Library) Manual.** Includes setup instructions, implementation details, and a function glossary.

## Acknowelgements

This work was possible through the generous support of NIH funding, Anderson and Robertson-Tessi acknowledge NCI U54CA193489, Anderson and Bravo acknowledge NCI UH2CA203781.

## Notes

#### Summary of Updates

a small correction to the manual regarding how PDEs are computed

