## Supplemental Figure 1: HAL Manual for "Hybrid Automata Library: A modular platform for efficient hybrid modeling with real-time visualization"

### HAL: Hybrid Automata Library

#### User Manual

**Abstract**

The Hybrid Automata Library (HAL) is a Java Library developed for use in mathematical oncology modeling. It is made of simple, efficient, generic components that can be used to model complex spatial systems. HAL's components can broadly be classified into: on- and off-lattice agent containers, finite difference diffusion fields, a GUI building system, and additional tools and utilities for computation and data collection. These components are designed to operate independently and are standardized to make them easy to interface with one another. As a demonstration of how modeling can be simplified using our approach, we have included a complete example of a hybrid model (a spatial model with interacting agent-based and PDE components). HAL is a useful asset for researchers who wish to build efficient 1D, 2D and 3D hybrid models in Java, while not starting entirely from scratch. It is available on github at https://github.com/MathOnco/HAL under the MIT License. HAL requires at least Java 8 or later to run, and the Java JDK version 1.8 or later to compile the source code.

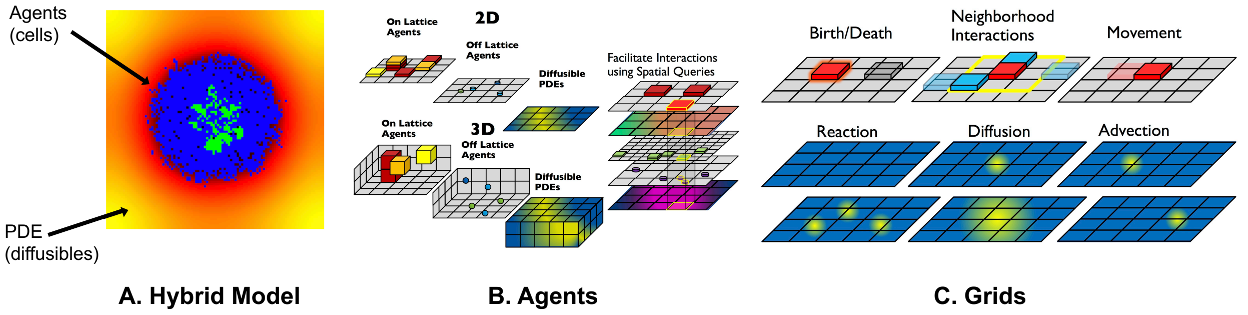

Figure 1: What is hybrid modeling? (A) A typical hybrid model consists of two types of components: agents (e.g tumor cells) and diffusibles described by partial differential equations. (B) HAL allows for agents to be off-lattice (able to move continously), on-lattice (bound to lattice positions) and can represent agents and PDEs in 1,2 and 3 spatial dimensions. Different agents and PDEs can interact spatially to construct complex models. (C) Agents are contained in grids, and have base actions of Birth/Death, interacting with neighbors, and movement. PDEs have base actions of Reaction (production and consumption of concentration) Diffusion (spreading of concentration) and Advection (directional flow of concentration)

### 1 Overview

Hybrid Automata Library (HAL) is a Java library that facilitates hybrid modeling: spatial models with interacting agent-based and partial-differential equation components. HAL's components can be broadly classified into: agent containers (on and off-lattice), finite difference diffusion fields, graphical user interface (GUI) components, and additional tools or utilities for computation and data manipulation. These components have a standardized interface that expedites the construction, analysis, and visualization of complex models.

HAL was originally developed to support mathematical oncology modeling efforts at the H. Lee Moffitt Cancer Center and Research Institute. To view several examples of projects built in HAL, since its inception in 2017, we direct the reader to the following website: halloworld.org. More details on the philosophy and technical details behind HAL can be found in the preprint on BioRxiv. The code base for HAL is maintained on GitHub, found here.

#### 1.1 What is hybrid modeling?

Hybrid modeling (see figure 1) is the integration of Agent-Based modeling and partial differential equation (PDE) modeling. It is commonly used in mathematical oncology to mechanistically model interactions between microenvironmental diffusibles (e.g drugs or resources) and agents (tumor cells). Tissue is represented using agent-based modeling, where each agent acts as a single cell in two- or three-dimensional space. As seen in figure 1, agents may be on-lattice or off-lattice, and exist in 2D or 3D (1D and 0D are also included). Agents are contained in grids. A single model may have multiple overlapping and interacting grids, representing moving and interacting cells, alongside diffusing drug and resources. Diffusibles that interact with the tissue are represented using partial differential equations (PDEs).

#### 1.2 Modularity

Each component (grids, agents) of HAL can function independently. This permits any combination of components to be used in a single model, with the use of spatial queries to combine them.

#### 1.3 Extensibility

HAL was designed to allow models and components to be extended and modified. Grids and agents from published models can be used as as a scaffold on which to do additional studies while keeping the prior work and their additions separated.

#### 1.4 Simplicity

Components are simple and generic making them applicable to a wide variety of modeling problems outside of mathematical oncology. A defensive programming paradigm was used to generate useful error messages when a component is used incorrectly. The purpose of this manual is to explain the modeling paradigm behind HAL, where the clear, consistent interface and methodology allows for ease of learning and implementation.

#### 1.5 Performance

HAL prioritizes performance in its algorithmic implementation. HAL includes efficient PDE solving algorithms, efficient visualization using BufferedImages and OpenGL, and leverages Java's impressive performance for executing ABM logic. These performance considerations allow for real-time display and visualization of models with minimal lag.

### 2 Setup

#### 2.1 Getting Java

In order to run models built using HAL's code base, you'll need to download at least the Java8 JDK (Java Development Kit); newer JDKs should also work. To check if the JDK is already installed, open a Command Line (or terminal) prompt and enter:

```
java -version
javac -version
```

If both of these commands print 1.8 or newer, you'll be able to run HAL. If not, get the latest Java JDK by googling it or from the Oracle Website. (No need to download the demos and samples).

#### 2.2 Getting the Framework Source Code

To download HAL's source code, go to the GitHub repository to clone HAL, or click "download" get a zip file containing the source code along with the included examples. You'll need to include these source files in each project developed using HAL. Unzip this folder and drop the files into the desired location.

#### 2.3 Getting Intellij IDEA

Intellij Idea is a great IDE to edit and run Java code. It can be downloaded at jetbrains.com (a community edition is freely available). Students and academic faculty members can get the professional version free by applying at jetbrains.com/student.

#### 2.4 Setting up the Project

1. Download HAL
2. Open Intellij Idea
   (a) Navigate to the File menu and click New -> "Project from Existing Sources"
   (b) Navigate to the directory with the unzipped HAL Source code
3. Click "yes" for each step until asked for the Java JDK
   (a) Mac: navigate to "/Library/ Java/ JavaVirtualMachines/"
   (b) Windows: navigate to "C:\ Program Files\ Java\"
4. Once the setup is complete we will need to do one more step and add some libraries that allow for 2D and 3D OpenGL visualization:
   (a) Navigate to the File menu and click "Project Structure"
   (b) Click the "Libraries" tab
   (c) Use the minus button (-) to remove any pre-existing libraries
   (d) Click the plus button, and direct the file browser to the "Framework/lib" folder.
   (e) Click apply or OK

This will setup the classes, sources, and native library locations that OpenGL needs. To test if everything is working properly, navigate to the "Examples" folder in the side menu, double click on any of the classes to open the source code. Scroll through the code until you find the class called "public static void main" and click the green arrow directly to the left to Run.

We've also designed a custom dark color scheme for Intellj Idea, which can be installed by navigating to the File menu and clicking "Import Settings" and navigating to the "HalColorSchemes.jar" file in the top level directory of HAL's source code.

#### 2.5 Features of Intellij Idea

- Automatically importing classes at first code mention (or, right click on the class name to import).
- Debugging with the "debugger" (find an excellent tutorial on the Intellij debugger here).
  - HINT: while paused in the debugger right click on anything and click "evaluate expression."
- "Refactor" to rename variables, change all function signatures, etc. in one click (right click on some code and check out the "refactor" submenu).
- Use "find usages" and "Go To Declaration" to move fluidly around your code base and to see how its pieces connect together (right clicking on things will get you there, see "find usages" and the "go to" submenu).
- using Shift-F10 to quickly run the currently open file, and many other hotkeys to speed up your workflow.
- Tapping Shift twice allows you to search the entire codebase for any search term.
- Type "fori" , "iter" , "sout" or "psvm" then type "tab" to create a for loop, foreach loop, print statment, and main function respectively.

### 3 Introduction

#### 3.1 Learning Java

The following sections assume a basic working knowledge of the Java language and Object-Oriented programming (OOP) methodology. To learn more about Java, there are many excellent tutorials on Youtube, Code Academy, and Learn Java Online.

#### 3.2 Learning HAL

The best way to learn HAL is to browse the "Examples" or "LEARN_HERE" folders. The first contains several in depth full models and the second gives detail on implementation of specific components (agents, grids, etc). Alternatively, several tutorials are posted to YouTube:

- HAL Tutorial 1: Setup
- HAL Tutorial 2: On Lattice Model
- HAL Tutorial 3: Intellij Tips and On Lattice Model Continued
- HAL Tutorial 4: Off Lattice

#### 3.3 Source Code Organization

**Examples**

This folder contains models that utilize a full range of HAL's features, including off-lattice and on-lattice agents, PDE solvers, UI visualization components, and more.

**LEARN_HERE**

This folder contains smaller examples that highlight specific framework components with clarity.

**Framework**

This folder contains all the components of the source code of HAL. In general, these files should not be changed or altered. Descriptions of the 7 subsections contained within the framework folder are detailed below.

#### 3.4 Components of HAL

**GridsAndAgents** Contains all of the Agent and Grid types that the framework supports. There are 2D and 3D, stackable and unstackable, on-lattice and off-lattice agents and grids to contain them. The base classes that the grids and agents extend are also in this folder.

**Gui** Contains the main UIWindow class, as well as many component classes that can be added to the UI.

**Tools** Contains many useful tool classes, such as a FileIO wrapper, a genetic algorithm class, a multiwell experiment runner, etc. as well as a Util class that is a container of generic static methods. It also contains classes that are used internally by these tools.

**Interfaces** Contains interfaces that are used by the framework internally. It is useful to reference this folder if a framework function takes a function interface argument, or implements an interface.

**Lib** Contains outside libraries that have been integrated with the framework.

**Util** Utility functions such as color functions, array functions, cellular automata neighborhood functions, and math functions.

**Rand** Contains functions used to generates random numbers from various distributions.

**Testing** Contains unit tests for many of HAL's components

### 4 Grids and Agents

There are 10 base types of agents that are named according to how many spatial dimensions they occupy, and how they are allowed to move in space. The SQ and PT suffixes refer to whether the agents are imagined to exist as lattice bound squares/voxels that move discretely, or as as non-volumetric points in space that move continuously.

AgentGrids are used as spatial containers for agents (1, 2, 3 dimensional and 0 dimensional or non-spatial). Internally, AgentGrids are composed of two datastructures: an agent list for agent iteration, and an agent lattice for spatial queries (even off-lattice agents are stored on a lattice for quick access). The agent list can be shuffled at every iteration to randomize iteration order, and the list holds onto removed agents to facilitate object recycling.

Similarly, PDE Grids consist of either a 1D, 2D, or 3D lattice of concentrations. PDE grids contain functions that will solve reaction-advection-diffusion equations. Currently implemented PDE solution methods include:

- Forward Difference in time and 2nd order central difference in space Diffusion
- ADI Diffusion
- 1st order upwind finite difference advection for incompressible flows
- 1st order finite volume upwind advection for compressible flows
- Modification of values at single lattice positions to facilitate reaction with agents or other sources/sinks.

Most of these methods are flexible, allowing for variable diffusion rates and advection velocities as well as different boundary conditions such as periodic, Dirichlet, and zero-flux Neumann.

#### 4.1 Types of Grid

**AgentGrid0D** Holds nonspatial agents

**AgentGrid1D** Holds all 1D agents

**AgentGrid2D** Holds all 2D agents

**AgentGrid3D** Holds all 3D agents

**PDEGrid1D** facilitates modeling a single diffusible field in 1D

**PDEGrid2D** facilitates modeling a single diffusible field in 2D

**PDEGrid3D** facilitates modeling a single diffusible field in 3D

#### 4.2 Types of Agent

| Name | Spatial Dimension | Lattice-Bound | Stackable |
|---|---|---|---|
| Agent0D | 0 | N/A | N/A |
| AgentSQ1D | 1 | Yes | Yes |
| AgentSQ1Dunstackable | 1 | Yes | No |
| AgentPT1D | 1 | No | Yes |
| AgentSQ2D | 2 | Yes | Yes |
| AgentSQ2Dunstackable | 2 | Yes | No |
| AgentPT2D | 2 | No | Yes |
| AgentSQ3D | 3 | Yes | Yes |
| AgentSQ3Dunstackable | 3 | Yes | No |
| AgentPT3D | 3 | No | Yes |

Table 1: Here we summarize the attributes of each agent type. Spatial Dimension refers to how many dimensions the agents exist in, Lattice-Bound refers to whether agents are bound to discrete lattice positions or can move continuously. Stackable refers to whether multiple agents can exist at the same position.

#### 4.3 Grid and Agent Class Definition

For demonstration we will reuse portions of code from the CompetitiveReleaseModel example (see section 9). When developing your own custom AgentGrid or Agent classes, begin by "extending" the relevant AgentGrid or Agent classes (listed above in Section 3). This extension is done with the following syntax:

**Project specific AgentGrid definition syntax:**

```
public class ExampleModel extends AgentGrid2D<ExCell> {
```

**Project specific Agent class definition syntax:**

```
class ExampleCell extends AgentSQ2Dunstackable<ExampleModel> {
```

Note the <> after the base class name. In Java this is called a generic type argument. This is how we tell the Grid what kinds of Agent it will store, and how we tell the Agents what kind of AgentGrid will store them. It is used by the ExampleModel and ExampleCell to identify each other, so that their constituent functions can return the proper type, and access each other's variables and methods.

#### 4.4 Grid Constructors

In order to create a class that extends any of the AgentGrids, you must provide a constructor.

**ExampleGrid Constructor**

```
public ExampleGrid(int x, int y, Rand generator) {
    super(x, y, ExampleCell.class);
```

The first line declares the constructor and arguments, the second line calls super, which is required since our class extends a class with a constructor. Super is used to call the constructor of the base class. Into super we pass what the AgentGrid2D needs for initialization: an x and y dimension, which define the size of the grid, and ExampleCell.class, which is the class object of the ExampleCell. We pass the class object so that the ExampleGrid can create ExampleCells for us, as described in the next section.

#### 4.5 Agent Initialization Functions

Note: **Do not define a constructor for your Agent classes**.

The AgentGrid that houses the agent will act as a "factory" for agents, and produce them with the NewAgent() function. This will return an agent that is either newly constructed, or an agent that has died and is being recycled. This returning of dead agents for reuse as new ones allows the model to run without tasking the garbage collector with removing all of the dead agents. This will increase the speed and decrease the memory footprint of your model. Instead of a constructor you should define some sort of initialization for your agents, which you do directly after the call to NewAgent.

###### Agent Initialization

```
int RESISTANT = 0;
int SENSITIVE = 1;
for (int i = 0; i < totalCells; i++) {
    if (rng.Double() < resistantProb) {
        NewAgentSQ(tumorNeighborhood[i]).type = RESISTANT;
    } else {
        NewAgentSQ(tumorNeighborhood[i]).type = SENSITIVE;
    }
}
```

These lines of code come from the InitTumor function that the ExampleModel calls once at the beginning of a simulation. We use a random number generator and an if-else statement to decide whether to create a sensitive or resistant cell. since this type property is the only information individual cells store in this model, setting it is all that is needed to initialize a new cell. We call the NewAgentSQ() function, and pass in an index ("tumorNeighborhood" is an array of starting indices to setup the tumor) that marks where to place the new agent.

#### 4.6 Grid Indexing

There are 3 different ways to describe or index locations on framework grids: single indexing (find the "$i^{th}$" agent), square index (find the agent at grid point x, y) and point indexing (similar to square indexing, but for PT agents). Calling any of these functions will return NULL when no agent is present at that location.

##### 4.6.1 Single Indexing

Since the x dimension, y dimension, and possibly the z dimension values are Grid constants, every square or voxel can be uniquely identified with a single integer index. Functions using this kind of indexing typically end with the SQ (abbreviating Square) phrase at the end of the function name. Single indexing is done with the following mappings:

In 2D: $I(x, y) = x * yDim + y$

In 3D: $I(x, y, z) = x * yDim * zDim + y * zDim + z$

the agents/values in the grids are stored as a single array, so single indexing is actually the most efficient as it requires no conversion.

##### 4.6.2 Square Indexing

Similar to single indexing, square indexing uses a set of integers to refer to a specific square or voxel, as an (x,y) or an (x,y,z) set. Functions using this kind of indexing typically end with the SQ (abbreviating Square) phrase at the end of the function name.

##### 4.6.3 Point Indexing

Uses a set of double values, to define continuous coordinates. Functions using this kind of indexing typically end with the PT (abbreviating Point) phrase at the end of the function name. The integer flooring of a coordinate set corresponds to the Square or Voxel that contains the point.

#### 4.7 Typical Grid Loop

Next, an example Loop or Run function shows how to iterate through all agents in order to call "Step()" on each agent, which may contain functions related to birth, death, or mutations, for example.

```
public void Run(){
  for (int i = 0; i < runTicks; i++) {
    for (GOLAgent a : this) {
      a.Step();
    }
  }
}
```

The outer for loop counts the ticks, meaning that we will run for a total of runTicks steps. The inner for loop iterates over all the agents in the grid ("this" here is the grid that calls the Run function), and inside the loop we call that agent's Step function, which is defined in the GOLAgent class from the GameOfLife example.

#### 4.8 Grids containing Grid objects

Grids typically are containers of "agent" objects, but object Grids can be constructed to contain any arbitrary object including other grids. It may be useful to have a grid of grid objects when doing multiple stochastic simulations along with many other scenarios.

```
Grid2Dobject<ExampleGrid> GlobalGrid = new Grid2Dobject<>(10,10);
for (int x = 0; x < 10; x++) {
```

```
    for (int y = 0; y < 10; y++) {
        GlobalGrid.Set(x,y,new ExampleGrid(100,100,new Rand()));
    }
}
```

In the above example, GlobalGrid contains 10 rows and 10 columns of ExampleGrid grids, which in turn contain 100 rows and 100 columns of ExampleCell objects.

#### 4.9 Assigning a Grid as an attribute

HAL allows for multiple grids overlaying each other. To access one grid from a second grid, it's ideal to set the first grid as an attribute of the second grid, as follows:

```
public class SecondGrid extends AgentGrid2D<ExampleAgent> {
  final public ExampleGrid firstGrid;

  // constructor
  SecondGrid(int x, int y, ExampleGrid firstGrid) {
    super(x, y, ExampleAgent.class);
    this.examplefirst = firstGrid;
  }
}
```

In the above example, the constructor for the second grid requires an argument for the first grid. After instantiation of SecondGrid, we have access to the firstGrid as a member of SecondGrid

#### 4.10 Type Hierarchy

In order to navigate the source code and see the full set of functions and properties of the framework components, it is important to become familiar with the type hierarchy that the framework uses. Figure 2 summarizes this hierarchy for 2D agents.

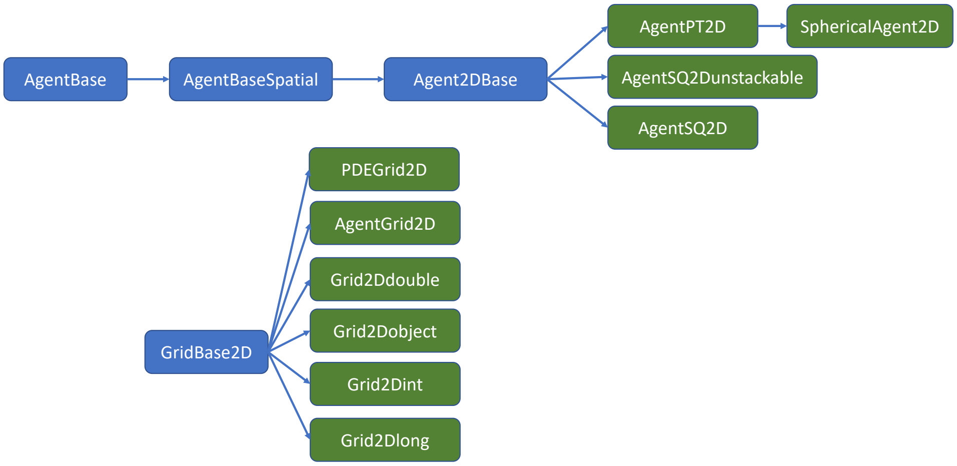

Figure 2:

Each node in the hierarchy names a class. each arrow denotes an extends relationship, eg. AgentBaseSpatial extends AgentBase. The blue classes are abstract and cannot be used directly. The green classes are fully implemented and can be used/extended further. Classes later in the hierarchy keep all properties and methods of the classes that they extend, which means that to see the full set of functions a class implements, one has to look at all of the extended classes beneath that class as well. The method summaries included in this manual can simplify this process somewhat as they show useful methods of the extended classes regardless of where they come from in the class hierarchy.

#### 4.11 Agent Functions and Properties

Here we describe the built-in Agent functions. (3D adds a z)

**G:** returns the grid that the agent belongs to (this is a permanent agent property, not a function)

**Age():** returns the age of the agent, in ticks. Be sure to use IncTick on the AgentGrid appropriately for this function to work.

**BirthTick():** returns the tick on which the agent was born

**SetBirthTick(tick):** sets the agent's birthTick value, which is used for the Age calculation

**IsAlive():** returns whether or not the agent currently exists on the grid

**Isq():** returns the index of the square that the agent is currently on

**Xsq(),Ysq():** returns the X or Y indices of the square that the agent is currently on.

**Xpt(),Ypt():** returns the X or Y coordinates of the agent. If the Agent is on-lattice, these functions will return the coordinates of the middle of the square that the agent is on.

**MoveSQ(x,y),MoveSQ(i):** moves the agent to the middle of the square at the indices/index specified

**MovePT(x,y):** moves the agent to the coordinates specified

**MoveSafeSQ(x,y),MoveSafePT(x,y):** Similar to the move functions, only it will automatically either apply wraparound (move the agent to the other side of the AgentGrid if it goes out of bounds and the Grid has enabled wraparound in that direction), or prevent moving along a partiular axis if movement would cause the agent to go out of bounds.

**Dispose():** removes the agent from the grid

**MapHood(int[]neighborhood):** This function takes a neighborhood (As generated by the neighborhood Util functions, or using GenHood2D/GenHood3D), and computes the set of position indices that the coordinates map to, with the agent at the $(0, 0)$ position of the neighborhood. The function returns the number of valid locations it found, which if wraparound is disabled may be less than the full set of neighborhood locations. See the CellStep function in the Complete Model example for more information.

**MapEmptyHood(int[]neighborhood),MapOccupiedHood(int[]neighborhood):** These functions are similar to the the MapHood function, but they will only include valid empty or occupied indices, and skip the others.

**MapHood(int[]neighborhood,CoordsToBool):** This function takes a neighborhood and another mapping function as argument. The mapping function should return a boolean that specifies whether coordinates map to a valid location. The function will only include valid indices, and skip the others.

**Xdisp(x),Ydisp(y):** returns the displacement from the agent to a given x or y

**Dist(x,y):** returns the distance from this agent to the given position

#### 4.12 SphericalAgent Functions and Properties

An extension of the AgentPT2D and AgentPT3D agent types, spherical agents have the additional property of a radius and x and y velocities. These properties allow spherical agents to behave like spheres that repel each other (3D adds a z).

**radius:** the radius property is used during SumForces to determine collisions

**xVel,yVel:** The x and y velocity properties are added to by SumForces and applied to the agent's position by calling ForceMove. Adding to these properties and calling ForceMove() will cause agents to move in a particular direction.

**Init(radius):** A default initialization function that sets the radius based on the argument, and the x and y velocities to 0.

**SumForces(interactionRad,OverlapForceResponse):** The interactionRad argument is a double that specifies how far apart to check for other agent centers to interact with, and should be set to the maximum distance apart that two interacting agent centers can be. The OverlapForceResponse argument must be a function that takes in an overlap and an agent, and returns a force response aka. (double,Agent) -> double. The double argument is the extent of the overlap. If this value is positive, then the two agents are "overlapping" by the distance specified. If the value is negative, then the two agents are separated by the distance specified.

The OverlapForceResponse should return a double which indicates the force to apply to the agent as a result of the overlap. If the force is positive, it will repel the agent away from the overlap direction, if it is negative it will pull it towards that direction. SumForces alters the xVel and yVel properites of the agent by calling OverlapForceResponse using every other agent within the interactionRad.

**ForceMove():** adds the xVel and yVel property values to the x,y position of the agent, moving it.

**Divide(divRadius,scratchCoordArr,Rand):** Facilitiates modeling cell division. The divRadius specifies how far apart from the center of the parent agent the daughters should be separated. The scratchCoordArr will store the randomly calculated axis of division. The axis is calculated using the Rand argument (HAL's random number generator object). If no Rand argument is provided, the values currently in the scratchCoordArr will be used to determine the axis of division. The first entry of scratchCoordArr is the x component of the axis, the second entry is the y component. Division is achieved by placing the newly generated daughter cell divRadius away from the parent cell using divCoordArr for the x and y components of the direction, and placing the parent divRadius away in the negative direction. Divide returns the newly created daughter cell.

**CapVelocity(maxVel):** caps the xVel and yVel variables such that their norm is not greater than maxVel

**ApplyFriction(frictionConst):** Multiplies xVel and yVel by frictionConst. If frictionConst = 1, then no friction force will be applied, if frictionConst = 0, then the cell's velocity components will be set to 0.

#### 4.13 Universal Grid Functions and Properties

These functions and properties are shared by all of the different types of grids (3D adds a z):

**xDim,yDim:** the x or y dimension of the Grid

**length:** the total number of squares in the grid, equivalent to xDim*yDim

**wrapX,wrapY:** boolean properties of the Grid that specify whether wraparound is enabled along the left and right edges of the Grid (wrapX) and the top and bottom edges of the Grid (wrapY).

**I(x,y):** converts a set of coordinates to the position index of the square at those coordinates.

**ItoX(i),ItoY(i):** converts an index of a square to that square's X or Y coordinate

**WrapI(x,y):** returns the index of the square at the provided x,y coordinates, with wraparound.

**In(x,y):** returns whether the x and y coordinates provided are inside the Grid.

**InWrap(x,y):** returns whether the x and y coordinates provided are inside the Grid, takes Grid-enabled wraparound into account.

**ConvXsq(x,otherGrid),ConvYsq(y,otherGrid):** returns the index of the center of the square in otherGrid that the coordinate maps to.

**ConvXpt(x,otherGrid),ConvYpt(y,otherGrid):** returns the position that the x/y argument rescales to in otherGrid

**DispX(x1,x2),DispY(y1,y2):** gets the displacement from the first coorinate to the second. if wraparound is enabled, the shortest displacement taking this into account will be returned.

**Dist(x1,y1,x2,y2):** gets the distance between two positions with or without grid wrap around (if wraparound is enabled, the shortest distance taking this into account will be returned)

**DistSquared(x1,y1,x2,y2):** gets the distance squared between two positions with or without grid wrap around (if wraparound is enabled, the shortest distance taking this into account will be returned). This is more efficient than the Dist function above as it skips a square-root calculation.

**MapHood(int[]hood,centerX,centerY):** This function takes a neighborhood array, translates the set of coordinates to be centered around a particular central location, and computes which indices the translated coordinates map to, putting the location indices in the hood array. The function returns the number of valid locations it set.

**MapHood(int[]hood,centerX,centerY,EvaluationFunction):** This function is very similar to the previous definition of MapHood, only it additionally takes as argument an EvaluationFunctoin. this function should take as argument (i,x,y) of a location and return a boolean that decides whether that location should be included as a valid one.

**IncTick():** increments the internal grid tick counter by 1. The grid tick is used with the Age() and BirthTick() functions to get age information about the agents on an AgentGrid. can otherwise be used as a counter with the other grid types.

**GetTick():** gets the current grid timestep.

**ResetTick():** sets the tick to 0.

**ApplyRectangle(startX,startY,width,height,ActionFunction):** applies the action function to all positions in the rectangle, will use wraparound if appropriate

**ApplyHood(int[]hood,centerX,centerY,ActionFunction):** applies the action function to all position in the neighborhood, centered around centerX,centerY. This function will ignore any neighborhood mapping, and will not modify any neighborhood mapping.

**ApplyHoodMapped(int[]hood,validCount,ActionFunction):** applies the action function to all positions in the neighborhood up to validCount, assumes the neighborhood is already mapped

**ContainsValidI(int[]hood,centerX,centerY,EvaluationFunction):** returns whether a valid index exists in the neighborhood

**BoundaryIs():** returns a list of indices, where each index maps to one square on the boundary of the grid

**AlongLineIs(x1,y1,x2,y2,int[]writeHere):** returns the set of indicies of squares that the line between (x1,y1) and (x2,y2) touches.

#### 4.14 AgentGrid Method Descriptions

Here we describe the builtin functions that the agent containing Grids expose to the user. Most functions that take x,y arguments can alternatively take a single index argument. The Grid will use the index to find the appropriate square (3D adds a z).

**NewAgentSQ(x,y),NewAgentSQ(i):** returns a new agent, which will be placed at the center of the indicated square. x,y, and i are assumed to be integers

**NewAgentPT(x,y):** returns a new agent, which will be placed at the coordinates specified. x and y are assumed to be doubles

**Pop():** returns the number of agents that are alive on the entire grid.

**CleanAgents():** reorders the list of agents so that dead agents will no longer have to be iterated over. Don't call this during the middle of iteration!

**ShuffleAgents():** shuffles the list of agents, so that they will no longer be iterated over in the same order. Don't call this during the middle of iteration!

**CleanShuffle():** Cleans the agent list and shuffles it. This function is often called at the end of a timestep. Don't call this during the middle of iteration!

**Reset():** disposes all agents, and resets the tick counter.

**MapHoodEmpty(int[]coords,centerX,centerY):** similar to the MapHood function, but will only include indices of locations that are empty

**MapHoodOccupied(int[]coords,centerX,centerY):** similar to the MapHood function, but will only include indices of locations that are occupied

**RandomAgent(rand):** returns a random living agent. Don't call this during the middle of iteration!

##### 4.14.1 AgentGrid Agent Search Functions

Several convenience methods have been added to make searching for agents easier.

**GetAgent(x,y)GetAgent(i):** Gets a single agent at the specified grid square, beware using this function with stackable agents, as it will only return one of the stack of agents. This function is recommended for the Unstackable Agents, as it tends to perform better than the other methods for single agent accesses.

**GetAgentSafe(x,y):** Same as GetAgent above, but if x or y are outside the domain, it will apply wrap around if wrapping is enabled, or return null.

**IterAgents(x,y),IterAgents(i):** use in a foreach loop to iterate over all agents at a location. example: for(AGENT_TYPE a : IterAgents(5,6)) will run a for loop over all agents at grid square (5,6) with the agent being stored as the "a" variable. be sure to set AGENT_TYPE to the type of agent that lives of the grid that you are iterating over.

**IterAgentsSafe(x,y):** Same as IterAgents above, but will apply wraparound if x,y fall outside the grid dimensions.

**IterAgentsRad(xPT,yPT,rad):** will iterate over all agents around the given coordinate pair that fall within radius rad.

**IterAgentsRadApprox(xPT,yPT,rad):** will iterate over all agents around the given coordinate pair that at least fall within radius rad, it is more efficient than IterAgentsRad, but it will also include some agents that fall outside rad, so be sure to do an additional distance check inside the function.

**IterAgentsHood(int[]hood,centerX,centerY):** will iterate over all agents in the neighborhood as though it were mapped to centerX,centerY position. Note that this function won't technically do the mapping. if the neighborhood is already mapped, use IterAgentsHoodMapped instead.

**IterAgentsHoodMapped(int[]hood,hoodLen):** will iterate over all agents in the already mapped neighborhood. Be sure to supply the proper hood length.

**IterAgentsRect(x,y,width,height):** iterates over all agents in the rectangle defined by (x,y) as the lower left corner, and (x+width,y+height) as the top right corner.

**GetAgents(ArrayList<AgentType>,x,y)GetAgentsHood...:** A variation of this function exists that matches every IterAgents variant mentioned above. Puts into the argument ArrayList all agents found at the specified location. Call the clear function on the ArrayList before passing it to GetAgents if you don't want to append to whatever agents were already added there.

**RandomAgent(x,y,rand,EvalAgent)RandomAgentHood...:** A variation of this function exists that matches every IterAgents variant mentioned above. These functions are used to query a location and choose a random agent from that area, an optional EvalAgent function (a function that takes an Agent and returns a boolean) can be provided to ensure that the random agent satisfies the condition

**PopAt(x,y),PopAt(i):** returns the number of agents that occupy the specified position. This is more efficient than using GetAgents and taking the length of the resulting ArrayList.

#### 4.15 PDEGrid Method Descriptions

The PDEGrid classes do not store agents. They are meant to be used to model Diffusible substances. PDEGrids contain two fields (arrays) of double values: a current field, and a next field. The intended usage of these fields is that the values from the current timestep are stored in the current field while changes due to transformations are accumulated in the next field. Central to the function of the PDEGrid is the Update() function, which updates the current field with the values in the next field. The functions included in the PDEGrid class are the following: (3D adds a z, 1D removes y).

Note: For the diffusion and advection schemes, the boundary is defined along the boundaries of the PDEGrid (0,0) to (xDim,yDim). In the case of ADI the boundary is defined at 1/2 a lattice position outside the edges of the usual lattice domain (for now...).

**Update():** adds the "next field" into the "current field". The values in the "current field" won't change unless Update() is called!

**Get(x,y):** returns the "current field" double array which is usually used as the main field that agents interact with

**Add(x,y,val):** adds the val to the "next field"

**Mul(x,y,val):** multiplies the val by the "current field" value at (x,y), and adds the product to the "next field"

**Set(x,y,val):** sets a value in the "next field". Any previous changes will be overwritten.

**MaxDifRecord():** returns the maximum difference in any single lattice position between the current field and the current field the last time MaxDifRecord was called. (the function saves a record of the state for future comparison whenever it is called)

**MaxDelta():** returns the maximum difference in any single lattice position between the current field and the next field.

**MaxDeltaScaled(denomOffset):** returns the maximum difference in any single lattice position between the current field and next field, but proportional to the current field value. the denom offset is added to the current field value to make sure that a divide by 0 error can't occur.

**Diffusion(diffCoef):** runs diffusion on the current field, adding the deltas to the next field. This form of the function assumes either a reflective or periodic (wraparound) boundary, depending on how the PDEGrid was specified. The diffCoef variable is the nondimensionalized diffusion conefficient. If the dimensionalized diffusion coefficient is $D$ then diffCoef can be found by computing $\frac{D*TimeStep}{SpaceStep^2}$ Note that if the diffCoef exceeds 0.25, this diffusion method will become numerically unstable.

**Diffusion(diffCoef,boundaryValue):** has the same effect as the above diffusion function without the boundary value argument, except rather than assuming zero flux, the boundary condition is set to either the boundaryValue, or wraparound, depending on how the PDEGrid was specified.

**DiffusionADI(diffCoef):** runs diffusion on the current field using the ADI (alternating direction implicit) method. Without a boundaryValue argument, a zero flux boundary condition is imposed. Wraparound will not work with ADI. ADI is numerically stable at any diffusion rate.

**DiffusionADI(diffCoef,boundaryValue):** runs diffusion on the current field using the ADI (alternating direction implicit) method. ADI is numerically stable at any diffusion rate. Adding a boundary value to the function call will cause boundary conditions to be imposed. for ADI boundary conditions, the boundary of the grid is defined at 1 lattice position outside the edges of the usual lattice domain

**Advection(xVel,yVel):** runs advection, which moves the concentrations using a constant flow with the x and y velocities passed. this signature of the function assumes wrap-around, so there can be no net flux of concentrations. Note that xVel and yVel are nondimensionalized velocity coefficients. If the dimensionalized advection velocity in the x direction is $V_x$ then xVel can be found by computing $\frac{V_x*TimeStep}{SpaceStep}$. If $|xVel| + |yVel| > 1$, the function will be unstable.

**Advection(xVel,yVel,bounaryValue):** runs advection as described above with a boundary value, meaning that the boundary value will advect in from the upwind direction, and the concentration will disappear in the downwind direction. If $|xVel| + |yVel| > 1$, the function will be unstable.

**GradientX(x,y),GradientY(x,y):** returns the gradient of the diffusible in a direction at the coordinates specified

**GetAvg():** returns the mean value of the field

**GetMax():** returns the max value in the field

**GetMin():** returns the min value in the field

**SetAll(val),SetAll(val[])AddAll(val)MulAll(val):** applies the Set/Add/Mul operations to all entries of the "current field", adding the deltas to the "next field"

#### 4.16 Griddouble/Gridint/Gridlong Method Descriptions

As an alternative to this class, it may be useful to simply employ a double/int/long array whose length is equal to the length of the other associated grids. The I() function of any associated grids can be used to access values in the double array with x,y or x,y,z coordinates.

**GetField():** returns the field double array that the grid stores

**Get(i),Get(x,y),Set(i),Set(x,y,val),Add(i),Add(x,y,val),Mul(i,val),Mul(x,y,val):** gets, sets, adds, or multiplies with a single value in the field at the specified coordinates

**SetAll(val),SetAll(val[])AddAll(val)MulAll(val):** applies the Set/Get/Mul operations to all entries of the field

**BoundAll(min,max),BoundAllSwap(min,max):** sets all values in the field so that they are between min and max

**GradientX(x,y),GradientY(x,y):** returns the gradient of the field in a direction at the coordinates specified

**GetAvg():** returns mean value of the grid

**GetMax():** returns the max value in the grid

**GetMin():** returns the min value in the grid

#### 4.17 AgentList Method Descriptions

AgentLists allow for keeping track of agent sub-populations. AgentLists must be instantiated with a type argument which specifies the type of agent that they will hold. When an agent dies, it is automatically removed from any AgentLists that contain it, so the AgentList will only contain living agents. It is not necissarily ordered. Use the java foreach loop syntax to iterate over the AgentList, just like the AgentGrids.

**AddAgent(agent):** adds an agent to the AgentList, agents can be added multiple times.

**RemoveAgent(agent):** removes all instances of the agent from the AgentList, returns the number of instances removed

**GetPop():** returns the size of the AgentList, duplicate agents will be counted multiple times

**InList(agent):** returns whether a given agent is already in the AgentList

**ShuffleAgents(rng):** Shuffles the agentlist order

**CleanAgents():** may speed up AgentList iteration if many agents from the AgentList have died recently

**RandomAgent(rng):** returns a random agent from the AgentList

### 5 Util.java

The Util class is one of the most ubiquitous classes in HAL. It contains all of the generic functions that wouldn't make sense to add to any particular object.

The list of utilities functions in HAL has only grown with time. The list presented here is not exhaustive, so I recommend looking at the file itself if you feel something is missing, on top of that feel free to ask for a new feature or send me an implementation and I'll most likely add it to the repository for everyone to share.

#### 5.1 Util Array Functions

A set of utilities for making array manipulation easier.

**ArrToString(arr,delim):** useful for collecting data or print statement debugging, returns the contents of an array as a single string. entries are separated by the delimeter argument

**IndicesArray(numEntries):** generates an array of ascending indices starting with 0, up to nEntries.

**Mean(arr):** returns the mean of an array

**Sum(arr):** returns the sum of an array

**Norm(arr):** returns the euclidean norm of an array

**NormSq(arr):** returns the squared euclidian norm of an array, somewhat more efficient than Norm

**SumTo1(arr):** scales all entries in an array so that their sum is 1.

**Normalize(arr):** scales all entries in an array so that their norm is 1.

#### 5.2 Util Neighborhood Functions

A set of utilities for generating neighborhoods. Neighborhoods are lists of x,y index pairs, of the form $[0_1, ...0_n, x_1, y_1, ...x_n, y_n]$ in the third dimension these are $[0_1, 0_2, ...0_n, x_1, y_1 z_1, ...x_n, y_n, z_n]$ these arrays are useful when finding the indices of the locations that make up the neighborhood around an agent when using Grid/Agent functions such as MapHood. Neighborhood arrays are always padded with 0s at the beginning, for use with the MapHood functions.

These functions should not be called over and over, as this would wastefully create arrays over and over. instead the function should be called once and be stored by the grid.

**GenHood2D(int[]coords),GenHood3D(int[]coords):** generates a 2D/3D neighborhood array for use with the MapHood functions. input should be an integer array of the form $[x_1, y_1, ...x_n, y_n]$ in 2D, or $[x_1, y_1 z_1, ...x_n, y_n, z_n]$ in 3D.

**VonNeumannHood(origin?),MooreHood(origin?):** returns an array that contains the coordinates of the VonNeumann or Moore Neigbhorhood in 2D. The boolean argument specifies whether the center of these neighborhood (0,0) should be included as part of the set of coordinates.

**VonNeumannHood3D(origin?),MooreHood3D(origin?):** returns an array that contains the coordinates of the VonNeumann or Moore Neigbhorhood in 3D. The boolean argument specifies whether the center of these neighborhood (0,0) should be included as part of the set of coordinates.

**HexHoodEvenY(origin?),HexHoodOddY(origin?):** to simulate a hex lattice on a Grid2D, the neighborhood used for a given agent changes depending on the position of the agent. The hex hood changes depending on whether the agent is on an even or odd Y position. These functions return an array that contains the proper coordinates for a given case.

**TriangleHoodSameParity(origin?),TriangleHoodDifParity(origin?):** to simulate a triangle lattice on a Grid2D, the neighborhood used for a given agent changes depending on the position of the agent. The Triangle hood changes depending on whether the parity ("even-ness") of the agents X and Y position match. These functions return an array that contains the proper coordinates for a given case.

**CircleHood(origin?,radius):** generates a neighborhood of all squares within the radius of a starting position at the middle of the (0,0) origin square. the boolean argument specifies whether (0,0) should be included in the return array.

**RectangleHood(origin?,radX,radY):** generates a neighborhood of all squares whose x displacement is within radX, and whose y displacement is within radY of a starting position at the middle of the (0,0) origin square. The boolean argument specifies whether (0,0) should be included in the return array.

**AlongLineCoords(x1,y1,x2,y2):** returns an array of all squares that touch a line between the two starting positions.

#### 5.3 Util Math Functions

**InfiniteLinesIntersection2D(x1,y1,x2,y2,x3,y3,x4,y4,double[]ret):** computes the intersection of lines between points 1 and 2, and points 3 and 4. puts the coordinates of the intersection point in ret. returns true if the infinite lines intersect (if they are not parallel). returns false if the lines are parallel.

**Bound(val,min,max):** returns the value bounded by the min and max.

**Rescale(val,min,max):** assumes the starting value is in the 0-1 scale, and rescales it be in the min-max scale.

**ModWrap(val,max):** wraps the value provided so that it must be between 0 and max. used to implement wraparound by the framework.

**ProtonsToPh(protonConc):** converts proton concentration to ph

**PhToProtons(ph):** converts ph to proton concentration

**ProbScale(prob,duration):** converts the probability that an even happens in unit time to the probability that that same event happens in duration time.

**Sigmoid(val,stretch,inflectionValue,minCap,maxCap):** val is the value that the sigmoid function is being applied to, the stretch argument stretchs or shrinks the sigmoid function along the x axis, the infectionValue governs where the inflection point of the sigmoid is, minCap and maxCap bound the sigmoid along the y axis.

#### 5.4 Util Misc Functions

**TimeStamp():** returns a timestamp string with format "YYYY_MM_DD_HH_MM_SS"

**PWD():** returns the current working directory as a string

**MemoryUsageStr():** returns a string with information about the current memory usage of the program.

**QuickSort(sortMe,greatestToLeast?):** requires the passed sortMe class to implement the Sortable interface. the passed boolean indicates whether the array should be sorted from greatest to least or least to greatest.

#### 5.5 Util Color Functions

These functions generate integers that store RGBA (Red,Green,Blue,Alpha) color channels internally, 8 bits per channel (integer values 0-255). These so-called "ColorInts" are intended to be used as arguments for the UIGrid, GridWindow, Vis2DOpenGL, and Vis3DOpenGL to set the color of the pixels or objects being displayed. The RGB components set the color, and the alpha component adds transparancy.

Note: All of these functions return a new ColorInt, as integers in java cannot be directly modified.

**RGB(r,g,b):** sets the rgb color channels using the continous 0-1 range mapping. The alpha is always set to 1.

**RGBA(r,g,b,a):** sets the rgb and alpha channels using the continuous 0-1 range mapping.

**RGB256(r,g,b):** sets the rgb color channels using the discrete 0-255 range mapping. The alpha is always set to 1.

**RGBA256(r,g,b,a):** sets the rgb and alpha channels using the 0-255 range mapping.

**GetRed(color),GetBlue(color),GetGreen(color),GetAlpha(color):** returns the value of a single channel using the continous 0-1 range mapping

**GetRed256(color),GetBlue256(color),GetGreen256(color),GetAlpha256(color):** returns the value of a single channel using the discrete 0-255 range mapping

**SetRed(color,r),SetGreen(color,g),SetBlue(color,b),SetAlpha(color,a):** returns a new ColorInt with the one of its channels changed to the argument passed in, uses the continuous 0-1 range mapping

**SetRed256(color,r),SetGreen256(color,g),SetBlue256(color,b),SetAlpha256(color,a):** returns a new ColorInt with one of its channels changed to the argument passed in, uses the discrete 0-255 range mapping

**CategoricalColor(index):** returns a categorical color from a nice mutually distinct set. Valid indices are 0-19. the color order is (blue,red,green,yellow,purple,orange,cyan,pink,brown,light blue,light red,light green,light yellow,light purple, light orange, light cyan,light pink, light brown, light gray, dark gray)

**HeatMapRGB(val),HeatMap???(val):** returns a new ColorInt using the heatmap color scale. Values are distinguished in the 0-1 range. the heatmap color scale is black at 0, white at 1, and transitions between these by changing one color channel at a time from none to full. The order that the channels are changed is dictated by the order of the 3 letters at the end of the function name. the possible orders are RGB, RBG, GRB, GBR, GRB and GBR

**HeatMapRGB(val,min,max),HeatMap???(val,min,max):** same as HeatMapRGB with more args; distinguishes values in the min-max range.

**HSBColor(hue,saturation,brightness):** returns a new colorInt using the HSB colorspace. Values are distinguished in the continuous 0-1 range.

**YCbCrColor(y,cb,cr):** returns a new colorInt using the YCbCr colorspace. Values are distinguished in the continous 0-1 range.

**CbCrPlaneColor(x,y):** returns a new colorInt using the CbCr plane at Y=0.5. A very nice colormap for distinguishing position in a 2 dimensional space. x,y are expected to be in the continuous 0-1 range.

#### 5.6 Util MultiThread Function

**Multithread(nRuns,nThreads,RunFun):** A function so useful that it deserves its own section, the multithread function creates a thread pool and launches a total of nRun threads, with nThreads running simultaneously at a time. The RunFun that is passed in must be a void function that takes an integer argument. When the function is called, this integer will be the index of that particular run in the lineup. This can be used to assign the result of many runs to a single array, for example, if the array is written to once by each RunFun at its run index. If you want to run many simulations simultaneously, this function is for you. Check out the MultithreadExample to see how to use this function.

#### 5.7 Util Save and Load Functions

**NOTE:SerializableModel_Interface** In order to save and load models, you must first have the model extend the SerializableModel interface (this interface is defined in the Interfaces folder). This interface has one method, called SetupConstructors. All you have to do in this method is call the AgentGrid function _PassAgentConstructor(AGENT.cass) once for each AgentGrid in your model, where AGENT is the type of agent that the AgentGrid holds. If your model does not use any AgentGrids, then this function can be left empty. See the LEARN_HERE/Agents/SaveLoadModel for an example.

**SaveState(model):** Saves a model state to a byte array and returns it. The model must implement the SerializableModel interface

**SaveState(model,fileName):** Saves a model state to a file with the name specified. Creates a new file or overwrites one if the file already exists. The model must implement the SerializableModel interface

**LoadState(byte[]state):** Loads a model form a byte array created with SaveState. The model must implement the SerializableModel interface

**LoadState(fileName):** Loads a model from a file created with SaveState. The model must implement the SerializableModel interface

### 6 Rand.java

**Int(bound):** generates an integer in the uniformly distributed range 0 up to but not including the bound value.

**Double():** generates a double in the range 0 to 1

**Double(bound):** generates a double in the uniformly distributed range 0 up to the bound value.

**Long(bound):** generates a long in the uniformly distributed range 0 up to but not including the bound value.

**Bool():** generates a random boolean value, with equal probability of true and false.

**Binomial(n,p):** samples the binomial distribution, returns the number of heads with n weighted coin flips and probability p of heads

**Multinomial(double[]probs,n,Binomial,int[]ret):** fills the return array with the number of occurrences of each event. n is the total number of occurrences to bin.

**Shuffle(arr,sampleSize,numberOfShuffles):** shuffles an array, sampleSize is how much of the complete array should be involved in shuffling, and numberOfShuffles is the number of entries of the array that will be shuffled. the shuffling results will always start from the beginning of the array up to numberOfShuffles.

**Gaussian(mean,stdDev):** samples a gaussian with the mean and standard deviation given.

**RandomVariable(double[]probs):** samples the distribution of probabilities (which should sum to 1, the SumTo1 function comes in handy here) and returns the index of the probability bin that was randomly chosen.

**RandomPointOnSphereEdge(radius,double[]ret):** writes into ret the coordinates of a random point on a sphere with given radius centered on (0,0,0)

**RandomPointInSphere(radius,double[]ret):** writes into ret the coordinates of a random point of inside a sphere with given radius centered on (0,0,0)

**RandomPointOnCircleEdge(radius,double[]ret):** writes into ret the coordinates of a random point on the edge of a circle with given radius centered on (0,0)

**RandomPointInCircle(radius,double[]ret):** writes into ret the coordinates of a random point on the inside a circle with given radius centered on (0,0)

### 7 Gui

What fun is a model without being able to see and play with it in real time? The Gui classes allow you to easily do this and works on top of the Java Swing gui system (except the Vis2DOpenGL and Vis3DOpenGL, which are built on lwjgl).

#### 7.1 Types of Gui and Method Descriptions

Here we list the different guis that are provided by HAL and provide a summary of their functions

##### 7.1.1 GridWindow

The simplest built-in Gui, it is nothing more than a UIGrid imbedded in a UIWindow. Recommended for first-time users. All functions described below after Close() are also shared with the UIGrid class

**GridWindow(title,xDim,yDim,scaleFactor,main?,active?):** sets up a GridWindow, the pixel dimensions of the created window will be: $Width = xDim * scaleFactor$ and $Height = yDim * scaleFactor$ the main boolean specifies whether the program should exit when the window is closed. the active boolean allows easily toggling the objects on and off.

**TickPause(milliseconds):** call this once per step of your model, and the function will ensure that your model runs at the rate provided in milliseconds. the function will take the amount time between calls into account to ensure a consistent tick rate.

**IsKeyDown(char):** returns whether the given key is currently pressed

**AddKeyResponse(OnKeyDown,OnKeyUp):** takes 2 key response functions that will be called whenever a key is pressed or released

**Close():** disposes of the GridWindow.

**Clear(color):** sets all pixels to a single color.

**SetPix(x,y,color),SetPix(i,color):** sets an individual pixel on the GridWindow. in the visualization the pixel will take up scaleFactor*scaleFactor screen pixels.

**SetPix(x,y,ColorIntGenerator),SetPix(i,ColorIntGenerator):** same functionality as SetPix with a color argument, but instead takes a ColorIntGenerator function (a function that takes no arguments and returns an int). The reason to use this method is that when the gui is inactivated the ColorIntGenerator function will not be called, which saves the computation time of generating the color.

**SetString(string,xLeft,yTop,color,bkColor):** draws a string to the GridWindow. The characters in the string have length 3 pixels, and height 5 pixels, with one pixel width between characters. All characters will be drawn to the same line.

**GetPix(x,y),GetPix(i):** returns the pixel color at that location as a colorInt

**PlotSegment(x1,y1,x2,y2,color,scaleX,scaleY):** Plots a line segment, connecting all pixels between the points defined by $(x1, y1)$ and $(x2, y2)$ with the provided color. If you are using this function on a per-timestep basis, I recommend setting individual pixels with SetPix, as it is more performant. The scaling variables adjust the spatial scale of the points.

**PlotLine(double[]xs,double[]ys,color,startPoint,endPoint,scaleX,scaleY):** plots a line by drawing segments between consecutive points. point $i$ is defined by $(xs[i], ys[i])$. Points are drawn starting at index startPoint, and ending at index endPoint. The scaling variables adjust the spatial scale of the points.

**PlotLine(double[]xys,color,startPoint,endPoint,scaleX,scaleY):** plots a line by drawing segments between consecutive points. Points are expected in the coords format $(x1, y1, x2, y2, x3, y3...)$. Point $i$ is defined by $(xys[i*2], xys[i*2+1])$. Points are drawn starting at index startPoint, and ending at index endPoint. The scaling variables adjust the spatial scale of the points.

**DrawStringSingleLine(string,xLeft,yTop,color,bkColor):** draws a string onto the GuiWindow.

**AddAlphaGrid(overlay):** adds another UIGrid as an overlay to compose with the main UIGrid. Alpha blending will be used to combine them.

##### 7.1.2 UIWindow

A container for Gui Components, which will be detailed below. The most supported of the gui types.

**UIWindow(title,main?,CloseAction,active?):** the title string is displayed in the top bar, the main boolean specifies whether the program should exit when the window is closed. The active boolean allows easily toggling the objects on and off.

**AddCol(column,component):** components are added to the gui from top to bottom in columns. When the window is displayed, the rows and columns expand to fit the largest element in each.

**RunGui():** once all components have been added, the RunGui function runs the gui and displays it to the screen.

**GetBool(label):** attempts to pull a boolean from the Param with the cooresponding label, works with the UIBoolInput

**GetInt(label):** attempts to pull an integer from the Param with the cooresponding label, works with the UIIntInput and UIComboBoxInput (returns the index of the chosen option)

**GetDouble(label):** attempts to pull a double from the Param with the cooresponding label, works with the UIIntInput and UIDoubleInput

**GetString(label):** attempts to pull a string from the Param with the cooresponding label, works with all Param types

**Dispose():** disposes of the GridWindow.

**TickPause(millis):** call this once per step of your model, and the function will ensure that your model runs at the rate provided in millis. The function will take the amount time between calls into account to ensure a consistent tick rate.

**IsKeyDown(char):** returns whether the given key is currently pressed

**AddKeyResponse(OnKeyDown,OnKeyUp):** takes 2 key response functions that will be called whenever a key is pressed or released

##### 7.1.3 OpenGL2DWindow

A window for visualizing 2D models, especially off lattice ones. I usually recommend using a UIGrid instead for 2D models.

**OpenGL2DWindow(xPix,yPix,xDim,yDim,title,active?):** creates a Vis2DOpenGL window. The dimensions on screen are xPix by yPix. the xDim and yDim dimensions should match the model being drawn. The title string is displayed on the top of the window, the active boolean allows easily disabling the Vis2DOpenGL

**Update():** renders all draw commands to the window

**IsClosed():** returns true if the close button has been clicked in the Gui

**Close():** closes the Vis2DOpenGL. Happens automatically when the main function finishes.

**Clear(clearColor):** usually called first, sets the screen to a color.

**Circle(x,y,z,radius,color):** Draws a circle. Currently this is the only builtin draw functionality along with the FanShape function from which it is derived.

**Line(x1,y1,x2,y2,color):** Draws a line between 2 points

**LineStrip(double[]xs,double[]ys,color):** draws a set of connected line segments, xs and ys are expected to be the same length.

**LineStrip(double[]coords,color):** draws a set of connected line segments, coords is expected to consist of [x1,y1,x2,y2...] pairs of point coordinates.

**TickPause(millis):** call this once per step of your model, and the function will ensure that your model runs at the rate provided in millis. The function will take the amount time between calls into account to ensure a consistent tick rate.

##### 7.1.4 OpenGL3DWindow

A window for visualizing 3D models.

**Vis3DOpenGL(xPix,yPix,xDim,yDim,title,active?):** creates a Vis3DOpenGL window. The dimensions on screen are xPix by yPix. The xDim, yDim, and zDim dimensions should match the model being drawn. The title string is displayed on the top of the window, the active boolean allows easily disabling the Vis3DOpenGL

**CONTROLS:** to move around inside an active Vis3DOpenGL window, click on the window, after which the following controls apply:

- **Esc Key:** Get the mouse back
- **Mouse Move:** Change look direction
- **W Key:** Move forward
- **S Key:** Move backward
- **D Key:** Move right

- **A Key:** Move left
- **Shift Key:** Move up
- **Space Key:** Move down
- **Q Key:** Temporarily increase move speed
- **E Key:** Temporarily decrease move speed

**Update():** renders all draw commands to the window

**Clear(color):** usually called before anything else: clears the gui

**CheckClosed():** returns true if the close button has been clicked in the Gui

**Close():** closes the Vis3DOpenGL. Happens automatically when the main function finishes.

**Circle(x,y,z,radius,color):** Draws a circle. Currently this is the only builtin draw functionality along with the FanShape function from which it is derived.

**CelSphere(x,y,z,radius,color):** Draws a cool looking cel-shaded sphere, which is really several Circle function calls in a row internally. Less performant than calling the Circle function.

**TickPause(millis):** call this once per step of your model, and the function will ensure that your model runs at the rate provided in millis. The function will take the amount time between calls into account to ensure a consistent tick rate.

**Line(x1,y1,z1,x2,y2,z2,color):** Draws a line between 2 points

**LineStrip(double[]xs,double[]ys,double[]zs,color):** draws a set of connected line segments, xs and ys are expected to be the same length.

**LineStrip(double[]coords,color):** draws a set of connected line segments, coords is expected to consist of [x1,y1,z1,x2,y2,z2...] triplets of point coordinates.

#### 7.2 Types of GuiComponent and Method Descriptions

##### 7.2.1 UIGrid

A grid of pixels that are each set individually. Very fast and useful for displaying the contents of grids

**UIGrid(gridW,gridH,scaleFactor,active?):** sets up a UIGrid, the pixel dimensions of the created area will be: $Width = xDim * scaleFactor$ and $Height = yDim * scaleFactor$. The active boolean allows easily toggling the objects on and off. The functions that the UIGrid can execute are included above. see the GridWindow for the list of UIGrid functions

##### 7.2.2 UILabel

A label that displays text on the Gui and can be continuously updated. The UILabel's on-screen size will remain fixed at whatever size is needed to render the string first passed to it.

###### 7.2.3 UIButton

A button that when clicked triggers an interrupting function

###### 7.2.4 UIBoolInput

A button that can be set and unset, must be labeled. Use the UIWindow Param functions to interact.

###### 7.2.5 UIIntInput

An input line that expects an integer, must be labeled. Use the UIWindow Param functions to interact.

###### 7.2.6 UIDoubleInput

An input line that expects a double, must be labeled. Use the UIWindow Param functions to interact.

###### 7.2.7 UIStringInput

An input line that takes any string, must be labeled. Use the UIWindow Param functions to interact.

###### 7.2.8 UIComboBoxInput

A dropdown menu of text options, must be labeled. Use the UIWindow Param functions to interact.

###### 7.2.9 UIFileChooserInput

A button that when clicked triggers a gui that facilitates choosing an existing file or creating one, must be labeled. Use the UIWindow Param functions to interact.

##### 7.3 GifMaker

The GifMaker object is used in combination with a UIGrid or GridWindow to make GIF videos of model runs.

**GifMaker(outputPath,timeBetweenFramesMS,loopContinuously?):** creates a GifMaker object. the outputPath argument specifies the name of the file that will be output. timeBettweenFramesMS specifies how many milliseconds the GIF should pause between frames. loopContinuously specifies whether the GIF should automatically restart when it finishes.

**AddFrame(UIGrid):** adds the current UIGrid state to the GIF as a single frame

**Close():** Close this GifMaker object

### 8 Tools

#### 8.1 Tools/ FileIO

An essential piece of HAL, the FileIO class facilitates easily writing to and reading from files. this is important for collecting data from your models as well as systematically paramaterizing them. the API for the FileIO object is discussed.

**FileIO(filename,mode):** the FileIO constructor expects a filename or path as a string, and a mode string, of which there are 6 options:

- **"r"** creates a FileIO in read mode, this FileIO is able to read text files
- **"w"** creates a FileIO in write mode, this FileIO is able to write to a new text file
- **"a"** creates a FileIO in append mode, this FileIO is able to append to an existing text file or write to a new file.
- **"rb"** creates a FileIO in read binary mode, this FileIO is able to read binary files
- **"wb"** creates a FileIO in write binary mode, this FileIO is able to write to binary files.
- **"ab"** creates a FileIO in append binary mode, this FileIO is able to append to an existing binary file or write to a new file.

The functions in the next sections are split up based on which mode was used to open the FileIO

**Close():** make sure to call this function when finished with the FileIO. The FileIO uses buffers internally to make writing more efficient. Without calling Close() the buffers may never be fully written out.

###### Read Mode Functions

**Read():** returns an ArrayList of Strings. each string is one line from the file

**ReadLine():** returns the next line from the file as a string

**ReadLineDelimit(delimeter),ReadLineIntDelimit(delimeter):** returns an array of strings,ints,or doubles, etc. Each entry is parsed using the delimeter.

**ReadDelimit(delimeter),ReadIntDelimit(delimeter):** returns an ArrayList of arrays of strings,ints,or doubles, etc. Each entry is parsed using the delimeter. Each array in the ArrayList contains data from one line of the file.

###### Write Mode Functions

**Write(string):** writes the string argument to a file

**WriteDelimit(arr,delimit):** writes the contents of the provided array to a file, entries are separated using the delimiter.

**ReadBinary Mode Functions**

**ReadBinBool(bool),ReadBinInt(int),ReadBinDouble(double):** read the next single value from the binary file.

**ReadBinBools(bool[]),ReadBinInts(int[]),ReadBinDoubles(double[]):** fills the array arugment with values read from the binary file.

**WriteBinary Mode Functions**

**WriteBinBool(bool),WriteBinInt(int),WriteBinDouble(double):** writes a single value to the binary file.

**WriteBinBools(bool[]),WriteBinInts(int[]),WriteBinDouble(double[]):** writes every entry in the array to the binary file.

#### 8.2 Tools/ MultiWellExperiment

the multiwell experiment class will visualize many models simultaneously on a single GridWindow. It can also run the models in parallel for better throughput.

**MultiWellExperiment(numWellsX,numWellsY,models[],visXdim,visYdim,borderColor,scaleFactor,StepFn,ColorFn):** the constructor for the MultiWellExperiment class. numWellsX, numWellsY define the number of well (model) rows, models[] is an array of the starting conditions of the models. visXdim, visYdim, scaleFactor define the x pixels, y pixels, and scaling of the visualization of each model. borderColor defines the color of the separator between models. StepFn is a function argument that takes a model, a well index, and a timestep as argument, and should update the model argument for one timestep. ColorFn is a function argument that takes a model, x, and y, and is used to set one pixel of the visualization.

**Run(numTicks,multiThread?,tickPause):** runs a multiwell experiment for a numTicks duration. if the multiThread boolean is set to true, the model execution will be multithreaded.

**Step():** runs a single timestep

**LoadWells(models[]):** sets up the multiwell experiment with a new array of models

**RunGif(numTicks,outFileName,recordPeriod,multithread?):** runs a multiwell experiment for a numTicks duration. If the multiThread boolean is set to true, the model execution will be multithreaded. Saves every recordPeriod fames to a gif.

#### 8.3 Tools/ Modularity/ ModuleSetManager

The ModuleSetManager class is used to store and use module objects. The type argument is the baseclass module type that the ModuleSetManager will manage. For a semi-complete example of the concept in action, see the ModuleSetExample code in the Examples folder.

**ModuleSetManager(baseClass):** the module base class object should define all of the method hooks that modules can use, the behavior of the base class object will be ignored

**AddModule(newModule):** adds a module to the ModuleSetManager. The module should override any functions in the baseClass that will be used with the model

**IterMethod(methodName):** use this function with a foreach loop to iterate over modules that override a given method. Used to run the module functions when appropriate

#### 8.4 Tools/ Modularity/ VarSetManager

The VarSetManager class maintains a count of double variables that are requested for an agent class that implements the VarSet interface. This tool is useful along with the ModuleSetManger to add agent variables that are only manipulated by one module. See the ModuleSetExample code in the Examples folder.

**NewVar():** generates a new variable as part of the var array, returns the index of the variable.

**AddVarSet(agent):** adds a new variable set to the given agent. does nothing if the variable set already exists for that agent.

#### 8.5 Tools/ PhylogenyTracker/ Genome

The genome class is useful for keeping track of shared genetic information and phylogenies. Extend this class to add any genetic/inherited information that needs to be tracked. An instance of the Genome class should be created once for every new mutation, and all agents with the same genome should share the same class instance as a member variable

**Genome(parent,removeLeaves?):** call this with the parent set to null to create a new phylogeny. The removeLeaves option specifies whether the phylogeny should continue to store dead leaves (lineages with no active individuals). This should also be called whenever a new clone is created, along with IncPop to add one individual to the new clone.

**GetId():** gets the ID of the genome, which indicates the order in which it arose

**GetPop():** returns the current active population that shares this genome

**GetParent():** returns the parent genome that was mutated to give rise to this genome

**SetPop(pop):** sets the active population size for this genome to a specific value.

**IncPop():** adds an individual to the genome population, should be called as part of the initialization of all agents that share this genome.

**DecPop():** removes an individual from the genome population, should be called as part of the disposal of all agents that share this genome.

**Traverse(GenomeFunction):** runs the GenomeFunction argument on every descendant of this genome

**TraverseWithLineage(ArrayList<Genome>lineageStorage,GenomeFunction):** runs the GenomeFunction argument on every descendant of this genome, will pass as argument the lineage from this genome to the descendant

**GetChildren(ArrayList<Genome>childrenStorage):** adds all direct descendants of the genome to the ArrayList

**GetLineage(ArrayList<Genome>lineageStorage):** adds all ancestors of the genome to the ArrayList

**GetNumGenomes():** returns the total number of genomes that have ever existed in the phylogeny

**GetNumLivingGenomes():** returns the number of currently active unique genomes

**GetNumTreeGenomes():** returns the number of genomes that exist in the phylogeny (not counting removed leaves)

**Pop():** returns the current population size that shares this genome

**PhylogenyPop():** returns the total population of all living members of the phylogeny

**GetRoot():** gets the first genome that started the phylogeny

### 9 Jar Files and Allocating Memory

Occasionally the user may benefit from increasing the maximum memory allocation pool for a Java virtual machine (JVM) upon execution. This can be accomplished using the "-Xmx" command with the proceeding memory allocation. For example if we wanted to specify 16 Gigabytes we would put -Xmx16G. This is particularly useful when running large-scale simulations on a high performance cluster (see jar file creation).

When executing a java jar file in command line use "java -Xmx16G -jar TargetJarFile.jar" to execute your model with extra memory. However, while in the IdeaJ environment this is accomplished by the following steps:

1. Click on "Edit Configurations" from the dropdown menu on the top right of Intellij next to the run button.
2. Specify the memory using the -Xmx argument in the "VM Arguments" box.

#### 9.1 Jar File Creation and Execution

When resources are needed that only a high-performance computing (HPC) environment can provide it may be useful to build a jar file of your model to execute independent of any IDE. IntelliJ IDEA makes it very easy to build this 'artifact.' There are two ways of executing your jar file. One is with command line arguments and the other is without. The process of building a jar file is the same for both, but with command line arguments you will set variables using 'args[].' With either case, the GUI should not be used within an HPC command line environment unless you are using port forwarding for graphics. This is dependent on your HPC environment and thus will not be covered here. To simply build a jar file from your project follow these steps:

1. Navigate to "File' -> 'Project Structure"
2. On the left, where we set up our libraries, click on "Artifacts"
3. Click the "+" button and select "JAR," finally click "From modules with dependencies...", You will only need to do this once!
4. Leave all the default options in place.
5. Click "Apply" and then "Ok"
6. From here go the top menu bar, click on "Build -> Build Artifacts..."

7. Using this you can select "Build," "Rebuild," "Clean", or edit which will take you back to Step 3.
8. Just choose "Build" (for the first time, you can use "Rebuild" later).
9. Your jar file is located within the project directory in the "out/artifacts/jarname/file.jar,"
10. the Jar file can be moved to another machine that has Java 8 or above for execution.
11. to run the jar file, enter something like: "Java -jar -Xmx16G /path/to/jar/file.jar arg1 arg2 arg3" where the optional flag "-Xmx16G" specifies that we want to use 16GB of memory for the program, "/path/to/jar/-file.jar" is the path to the jar file, and "arg1 arg2 arg3" are optional arguments that will show up in your program in the "String[] args" array in the main function.

### 10 Example: Competitive Release Model

To demonstrate how the aforementioned principles and components of HAL are applied, we now look at a simple but complete example of a hybrid modeling experiment. We will be designing a model of pulsed therapy based on a publication from our lab [1]. We also showcase the flexibility that the modular component approach brings by displaying 3 different parameterizations of the same model side by side in a "multiwell experiment".

#### 10.1 Competitive Release Introduction

As in [1], the presented model assumes two competeing tumor-cell phenotypes: a rapidly dividing, drug-sensitive phenotype and a slower dividing, drug-resistant phenotype. There is also a drug diffusible that enters the system through the model edges and is consumed over time by the tumor cells.

Every tick (timestep), each cell has a probability of death and a probability of division. The division probability is affected by phenotype and the availability of space. The death probability is affected by phenotype and the local drug concentration.

An interesting outcome of the experiment is that pulsed therapy is better at managing the tumor than constant therapy. The modular design of HAL allows us to test 3 different treatment conditions, each with an identical starting tumor (No drug, constant drug, and pulsed drug) Under pulsed therapy the sensitive population is kept in check, while still competing spatially with the resistant phenotype and preventing its expansion. The rest of the section describes in detail how this abstract model is generated.

Figure 3 provides a high level look at the structure of the code. Table **??** provides a quick reference for the built-in HAL functions in this example. Any functions that are used by the example but do not exist in the table are defined within the example itself and explained in detail below the code. Those fluent in Java may be able to understand the example just by reading the code and using the reference table.

Built-in framework functions and classes used in the code are highlighted in green to make identifying framework components easier

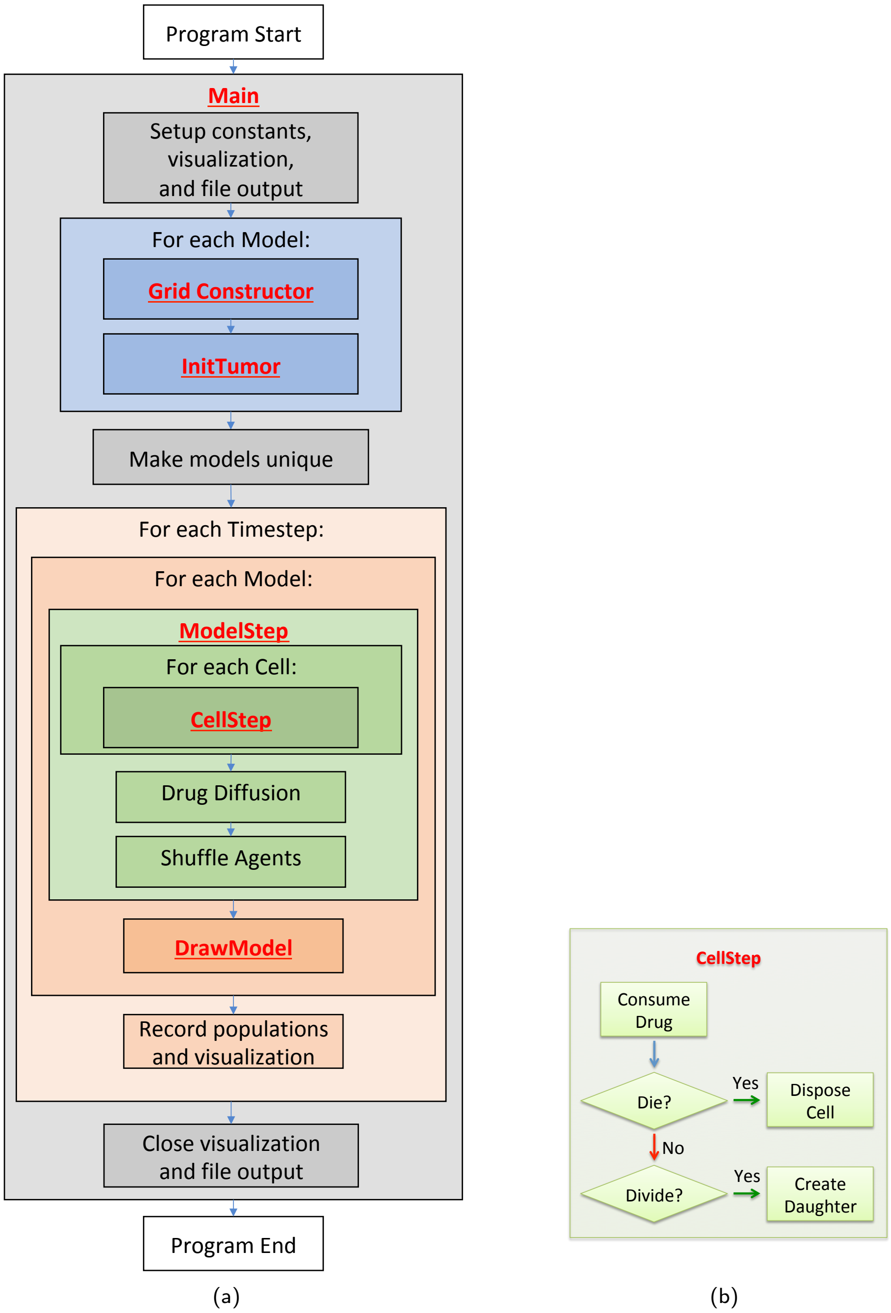

Figure 3: **(a)** Example program flow diagram. Red font indicates where functions are first called. **(b)** flow diagram of the CellStep function

| Function | Object | Action |
|---|---|---|
| NewAgentSQ(INDEX) | AgentGrid2D | Returns a new agent, placed at the center of the square at the provided position INDEX. |
| ShuffleAgents(RNG) | AgentGrid2D | Usually called after every timestep to shuffle the order of agent iteration. |
| GetTick() | AgentGrid2D | Returns the current grid timestep. |
| ItoX(INDEX), ItoY(INDEX) | AgentGrid2D | Converts from a grid position INDEX into the x and y components that point to the same grid position. |
| MapHood( NEIGHBORHOOD,X,Y) | AgentGrid2D | Finds all position indices in the provided neighborhood, centered around X,Y that don't fall out of bounds of the AgentGrid2D. Writes these indices into the NEIGHBORHOOD argument, and returns the number that were found. |
| MapEmptyHood( NEIGHBORHOOD) | AgentSQ2D | Finds all position indices in the provided neighborhood, centered around the agent, that do not have an agent occupying them. Writes these indices into the NEIGHBORHOOD argument, and returns the number that were found. |
| G | AgentSQ2D | Gets the grid that the agent occupies. |
| Isq() | AgentSQ2D | Gets the position index of the grid square that the agent occupies. |
| Dispose() | AgentSQ2D | Removes the agent from the grid and from iteration. |
| Get(INDEX) | PDEGrid2D | Returns the concentration of the PDE field at the given index. |
| Mul(INDEX, VALUE) | PDEGrid2D | Multiplies the concentration at the given INDEX by VALUE and adds the result to the current concentration when Update() is called |
| DiffusionADI( RATE) | PDEGrid2D | Applies diffusion using the ADI method with the rate constant provided. A reflective boundary is assumed. The result is applied when Update() is called. |
| DiffusionADI( RATE, BOUNDARY_COND) | PDEGrid2D | Applies diffusion using the ADI method with the RATE constant provided. The BOUNDARY_COND value diffuses from the grid borders. The result is applied when Update() is called. |
| Update() | PDEGrid2D | Applies all state changes simultaneously to the PDEGrid |
| SetPix(INDEX, COLOR) | GridWindow | Sets the color of a pixel. |
| TickPause( MILLISECONDS) | GridWindow | Pauses the program between calls to TickPause. The function automatically subtracts the time between calls from MILLISECONDS to ensure a consistent timestep rate for visualization. |
| ToPNG(FILENAME) | GridWindow | Writes out the current state of the UIWindow to a PNG image file. |
| Close() | GridWindow | Closes the GridWindow. |
| RGB(RED, GREEN, BLUE) | Util | Returns an integer with the requested color in RGB format. This value can be used for visualization. |
| HeatMapRGB(VALUE) | Util | Maps the VALUE argument (assumed to be between 0 and 1) to a color in the heat colormap. |
| CircleHood( INCLUDE_ORIGIN, RADIUS) | Util | Returns a set of coordinate pairs that define the neighborhood of all squares whose centers are within the RADIUS distance of the center $(0,0)$ origin square. The INCLUDE_ORIGIN argument specifies whether to include the origin in this set of coordinates. |
| MooreHood( INCLUDE_ORIGIN) | Util | Returns a set of coordinate pairs that define a Moore neighborhood around the $(0,0)$ origin square. The INCLUDE_ORIGIN boolean specifies whether we intend to include the origin in this set of coordinates. |
| Write(STRING) | FileIO | Writes the STRING to the output file. |
| Close() | FileIO | Closes the output file. |
| Double() | Rand | Generates a random double value in the range $[0-1)$ |

Table 2: HAL functions used in the example. Each function is a method of a particular object, meaning that when the function is called it may use properties that pertain to the object that it is called from.

#### 10.2 Main Function

We first examine the 'main' function for a bird's-eye view of how the program is structured. Source code elements highlighted in red are built-in HAL functions and objects, and can be referenced in Table **??**.

Note: Source code elements highlighted in red are already built into HAL.

```
1    public static void main(String[] args) {
2        //setting up starting constants and data collection
3        int x = 100, y = 100, visScale = 5, tumorRad = 10, msPause = 0;
4        double resistantProb = 0.5;
5        GridWindow win = new GridWindow("Competitive Release", x * 3, y, visScale);
6        FileIO popsOut = new FileIO("populations.csv", "w");
7        //setting up models
8        ExampleModel[] models = new ExampleModel[3];
9        for (int i = 0; i < models.length; i++) {
10           models[i] = new ExampleModel(x, y, new Rand());
11           models[i].InitTumor(tumorRad, resistantProb);
12       }
13       models[0].DRUG_DURATION = 0;//no drug
14       models[1].DRUG_DURATION = 200;//constant drug
15       //Main run loop
16       for (int tick = 0; tick < 10000; tick++) {
17           win.TickPause(msPause);
18           for (int i = 0; i < models.length; i++) {
19               models[i].ModelStep(tick);
20               models[i].DrawModel(win, i);
21           }
22           //data recording
23           popsOut.Write(models[0].Pop() + "," + models[1].Pop() + "," + models[2].Pop() +
                 "\n");
24           if ((tick) % 100 == 0) {
25               win.ToPNG("ModelsTick" + tick + ".png");
26           }
27       }
28       //closing data collection
29       popsOut.Close();
30       win.Close();
31   }
```

**Lines 3-4:** Defines all of the constants that will be needed to setup the model and display.

**5:** Creates a GridWindow of RGB pixels for visualization and for generating timestep PNG images. x*3, y define the dimensions of the pixel grid. the x variable is multiplied by 3 so that 3 models can be visualized side by side in the same window. The last argument is a scaling factor that specifies that each pixel on the grid will be viewed as a 5x5 square of pixels on the screen.

**6:** Creates a file output object that will write to a file called populations.csv.

**8:** Creates an array with 3 entries that will be populated with models.

**9-12:** Fills the model list with models that are initialized identically. Each model will hold and update its own cells and diffusible drug. See the Grid Definition and Constructor section and the InitTumor Function section for more details.

**13-14:** Setting the DRUG_DURATION constant creates the only difference in the 3 models being compared. In models[0] no drug will be administered. In models[1] drug administration will be constant (DRUG_DURATION is set equal to DRUG_CYCLE). In models[2] drug will be administered periodically (the default value of DRUG_DURATION is 40). See the ExampleModel Constructor and Properties section for the default model initialization.

**16:** Executes the main loop for 10000 timesteps.

**17:** Requires every iteration of the loop to take a minimum number of milliseconds. This slows down the execution and display of the model and makes it easier for the viewer to follow.

**18:** Loops over all models to update them.

**19:** Advances the state of the agents and diffusibles in each model by one timestep. See the Model Step Function for more details.

**20:** Draws the current state of each model to the window. See the Draw Model Function for more details.

**23:** Writes the population sizes of each model every timestep to allow the models to be compared.

**24:** Every 100 timesteps, writes the state of the model as captured by the GridWindow to a PNG file.

**29-30:** After the main for loop has finished, the FileIO object and the visualization window are closed, and the program ends.

#### 10.3 ExampleModel Constructor and Properties

This section explains how the grid is defined and instantiated.

```
1  public class ExampleModel extends AgentGrid2D<ExampleCell> {
2      //model constants
3      public final static int RESISTANT = RGB(0, 1, 0), SENSITIVE = RGB(0, 0, 1);
4      public double DIV_PROB_SEN = 0.025, DIV_PROB_RES = 0.01, DEATH_PROB = 0.001,
5              DRUG_DIFF_RATE = 2, DRUG_UPTAKE = -0.09, DRUG_TOXICITY = 0.2, DRUG_BOUNDARY_VAL
                   = 1.0;
6      public int DRUG_START = 400, DRUG_CYCLE = 200, DRUG_DURATION = 40;
7      //internal model objects
8      public PDEGrid2D drug;
9      public Rand rng;
10     public int[] divHood = MooreHood(false);
11
12     public ExampleModel(int x, int y, Rand generator) {
13         super(x, y, ExampleCell.class);
14         rng = generator;
15         drug = new PDEGrid2D(x, y);
16     }
```

This section covers how the grid is defined and instantiated.

**1:** The ExampleModel class, which is user defined and specific to this example, is built by extending the generic AgentGrid2D class. The extended grid class requires an agent type parameter, which specifies the type of agent that will live on the grid. To meet this requirement, the <ExampleCell> type parameter is added to the declaration.

**3:** Defines RESISTANT and SENSITIVE constants, which are created by the Util RGB function. These constants serve as both colors for drawing and as labels for the different cell types.

**4-7:** Defines all constants that will be needed during the model run. These values can be reassigned after model creation to test different parameter settings. In the main function, the DRUG_DURATION variable is modified for the Constant-Drug, and Pulsed Therapy experiment cases.

**9:** Declares that the model will contain a PDEGrid2D, which will hold the drug concentrations. The PDEGrid2D can only be initialized when the x and y dimensions of the model are known, which is why we do not assign their values until the constructor function is called.

**10:** Declares that the Grid will contain a Random number generator (the Rand object), but takes it in as a constructor argument to allow the modeler to seed the generator if desired for consistent output.

**11:** Creates a neighborhood using the MooreHood function. The MooreHood function generates a set of coordinates that define the Moore Neighborhood (the 8 closest coordinates to a central origin), centered around $(0, 0)$. The false argument declares that we do not want to include the origin in the neighborhood, just the 8 coordinates around that position. The neighborhood is stored in the format $[0_1 0_2, ..., 0_n, x_1, y_1, x_2, y_2, ..., x_n, y_n]$ . The leading zeros are written to when MapHood is called, and will store the position indices that the neighborhood maps to. See the CellStep function for more information, and the InitTumor Function Line 3 for another example of the use of neighborhoods

**13:** Defines the model constructor, which takes as arguments the x and y dimensions of the model and a random number generator (a Rand object).

**14:** Calls the AgentGrid2D constructor with super, passing it the x and y dimensions of the model, and the ExampleCell Class. This Class is used by the Grid to generate a new cell when the NewAgentSQ function is called.

**15-16:** The Rand argument is assigned and the drug PDEGrid2D is defined with matching dimensions.

#### 10.4 InitTumor Function

The next segment of code is a function from the ExampleModel class that defines how the tumor is first seeded after the ExampleModel is created.

```
1      public void InitTumor(double radius, double resistantProb) {
2          //get a list of indices that fill a circle at the center of the grid
3          int[] tumorNeighborhood = CircleHood(true, radius);
4          int hoodSize = MapHood(tumorNeighborhood, xDim / 2, yDim / 2);
5          for (int i = 0; i < hoodSize; i++) {
6              if (rng.Double() < resistantProb) {
7                  NewAgentSQ(tumorNeighborhood[i]).type = RESISTANT;
8              } else {
9                  NewAgentSQ(tumorNeighborhood[i]).type = SENSITIVE;
10             }
11         }
12     }
```

The next segment of code is a function from the ExampleModel class that defines how the tumor is first seeded after the ExampleModel is created.

**1:** The arguments passed to the InitTumor function are the approximate radius of the circular tumor being created and the probability that each created cell will be of the resistant phenotype.

**3:** Sets the tumorNeighborhood array using the CircleHood function, which stores coordinates in the form $[0_1, 0_2, ..., 0_n, x_1, y_1, x_2, y_2, ...x_n, y_n]$. The x,y coordinate pairs define a neighborhood of all squares whose centers are within the radius distance of the center $(0, 0)$ origin square. The leading 0s are used by the MapHood function to store the mapped indices. The boolean argument specifies that the origin will be included in this set of squares, thus making a completely filled circle of squares.

**4:** Uses the MapHood function to map the neighborhood defined above to be centered around xDim/2,yDim/2 (the dimensions of the AgentGrid). The results of the mapping are written as position indices to the beginning of the tumorNeighborhood array. MapHood returns the number of valid indices found, and this will be the size of the starting population.

**5:** Loops from 0 to hoodSize, allowing access to each mapped position index in the tumorNeighborhood.

**6:** Samples a random number in the range $[0 - 1)$ and compares to the resistantProb argument to set whether the cell should have the resistant phenotype or the sensitive phenotype.

**7-9:** Uses the NewAgentSQ function to place a new cell at each tumorNeighborhood position. In the same line we also specify that the phenotype should be either resistant or sensitive, depending on the result of the rng.Double() call.

#### 10.5 ModelStep Function

The next segment of code is a function from the ExampleModel class that defines how the tumor is first seeded after the ExampleModel is created.

```
1      public void ModelStep(int tick) {
2          ShuffleAgents(rng);
3          for (ExampleCell cell : this) {
4              cell.CellStep();
5          }
6          int periodTick = (tick - DRUG_START) % DRUG_CYCLE;
7          if (periodTick > 0 && periodTick < DRUG_DURATION) {
8              //drug will enter through boundaries
9              drug.DiffusionADI(DRUG_DIFF_RATE, DRUG_BOUNDARY_VAL);
10         } else {
11             //drug will not enter through boundaries
12             drug.DiffusionADI(DRUG_DIFF_RATE);
13         }
14     drug.Update()
15     }
```

This section looks at the main step function which is executed once per tick by the Model.

**2:** The ShuffleAgents function randomizes the order of iteration so that the agents are always looped through in random order.

**3-4:** Iterates over every cell on the grid, and calls the CellStep function on every cell.

**6:** The GetTick function is a built-in function that returns the current Grid tick. The If statement logic checks if the tick is past the drug start and if the tick is in the right part of the drug cycle to apply drug. (See the Grid Definition and Constructor section for the values of the constants involved, the DRUG_DURATION variable is set differently for each model in the Main Function)

**7-9:** If it is time to add drug to the model, the built-in DiffusionADI function is called. The default Diffusion function uses the standard 2D Laplacian and is of the form: $\frac{\delta C}{\delta t} = D\nabla^2 C$, where D in this case is the DRUG_DIFF_RATE. DiffusionADI uses the ADI method which is more stable and allows us to take larger steps than the 2D Laplacian can support. The additional argument to the DiffusionADI equation specifies the boundary condition value DRUG_BOUNDARY_VAL. This causes the drug to diffuse into the PDEGrid2D from the boundary.

**12:** Without the second argument the DiffusionADI function assumes a reflective boundary, meaning that drug concentration cannot escape or enter through the sides. Therefore the only way for the drug concentration to decrease is via consumption by the Cells. See the CellStep function section, line 6, for more information.

**14:** Update is called to apply the reaction and diffusion changes to th PDEGrid

#### 10.6 CellStep Function and Cell Properties

We next look at how the ExampleCell Agent is defined and at the CellStep function that runs once per Cell per timestep. The G property that is referenced many times in this section is a built-in agent property that gives access to the ExampleGrid object that the cell lives on.

```
1  class ExampleCell extends AgentSQ2Dunstackable<ExampleModel> {
2      public int type;
3
4      public void CellStep() {
5          //Consumption of Drug
6          G.drug.Mul(Isq(), G.DRUG_UPTAKE);
7          double deathProb, divProb;
8          //Chance of Death, depends on resistance and drug concentration
9          if (this.type == RESISTANT) {
10             deathProb = G.DEATH_PROB;
11         } else {
12             deathProb = G.DEATH_PROB + G.drug.Get(Isq()) * G.DRUG_TOXICITY;
13         }
14         if (G.rng.Double() < deathProb) {
15             Dispose();
16             return;
17         }
18         //Chance of Division, depends on resistance
19         if (this.type == RESISTANT) {
20             divProb = G.DIV_PROB_RES;
21         } else {
22             divProb = G.DIV_PROB_SEN;
23         }
24         if (G.rng.Double() < divProb) {
25             int options = MapEmptyHood(G.divHood);
26             if (options > 0) {
27                 G.NewAgentSQ(G.divHood[G.rng.Int(options)]).type = this.type;
28             }
29         }
```

```
30        }
31 }
```

**1:** The ExampleCell class is built by extending the generic AgentSQ2Dunstackable class. The extended Agent class requires the ExampleModel class as a type argument, which is the type of Grid that the Agent will live on. To meet this requirement, we add the <ExampleModel> type parameter to the extension.

**2:** Defines a cell property called "type". Each Cell holds a value for this field. If the value is RESISTANT, the Cell is of the resistant phenotype, if the value is SENSITIVE, the cell is of the sensitive phenotype. The RESISTANT and SENSITIVE values are defined in the ExampleGrid as constants (See the ExampleModel Constructor and Properties, line 3).

**6:** The G property is used to access the ExampleGrid object that the Cell lives on. G is used often with agent functions as the AgentGrid is expected to contain any information that is not local to the individual agent. Here it is used to get the drug PDEGrid2D. The drug concentration at the index that the Cell is currently occupying (Isq()) is then multiplied by the drug uptake constant, thus modeling local drug uptake by the Cell.

**7:** Defines deathProb and divProb variables, these will be assigned different values depending on whether the ExampleCell is RESISTANT or SENSITIVE.

**9-12:** If the cell is resistant, the deathProb variable is set to the DEATH_PROB value alone, if the cell is sensitive, an additional term is added to account for the probability of the cell dying from drug exposure, using the concentration of drug at the cell's position (Isq())

**14-16:** Samples a random number in the range $[0-1)$ and compares to deathProb to determine whether the cell will die. If so, the built-in agent Dispose() function is called, which removes the agent from the grid, and then return is called so that the dead cell will not divide.

**19-22:** Sets the divProb variable to either DIV_PROB_RES for resistant cells, or DIV_PROB_SEN for sensitive cells.

**24:** Samples a random number in the range $[0-1)$ and compares to divProb to determine whether the cell will divide.

**25:** If the cell divides, the MapEmptyHood function is used, which checks the positions in the divHood (the Moore neighborhood as defined in the ExampleModel Constructor and Properties section, line 11) around the Cell, and writes the position indices that do not contain any agents into the divHood. MapEmptyHood returns the number of valid empty positions found.

**26-27:** If there are one or more valid division options, a new daughter cell is created using the NewAgentSQ function and its starting location is chosen by randomly sampling the divHood array to pull out one if its valid locations. The other daughter is assumed to occupy the same location as the mother cell. Finally with the .type=this.type statement, the phenotype of the newly placed daughter cell is inherited from the mother cell.

#### 10.7 DrawModel Function

We next look at the DrawModel Function, which is used to display a summary of the model state on a GridWindow object. DrawModel is called once for each model per timestep, see the Main Function section for more information.

```
1     public void DrawModel(GridWindow vis, int iModel) {
2         for (int x = 0; x < xDim; x++) {
3             for (int y = 0; y < yDim; y++) {
4                 ExampleCell drawMe = GetAgent(x, y);
5                 if (drawMe != null) {
6                     vis.SetPix(x + iModel * xDim, y, drawMe.type);
7                 } else {
8                     vis.SetPix(x + iModel * xDim, y, HeatMapRGB(drug.Get(x, y)));
9                 }
10            }
11        }
12    }
```

**2-3:** Loops over every lattice position of the grid being drawn, xDim and yDim refer to the dimensions of the model.

**4:** Uses the GetAgent function to get the Cell that is at the x,y position.

**5-6:** If a cell exists at the requested position, the corresponding pixel on the GridWindow is set to the cell's phenotype color. To draw the models side by side, the pixel being drawn is displaced to the right by the model index.

**7-8:** If there is no cell to draw, then the pixel color is set based on the drug concentration at the same index, using the built-in heat colormap.

#### 10.8 Imports

The final code snippet looks at the imports that are needed. Any modern Java IDE should generate import statements automatically.

```
1  package Examples._6CompetitiveRelease;
2  import Framework.GridsAndAgents.AgentGrid2D;
3  import Framework.GridsAndAgents.PDEGrid2D;
4  import Framework.Gui.GridWindow;
5  import Framework.GridsAndAgents.AgentSQ2Dunstackable;
6  import Framework.Gui.UIGrid;
7  import Framework.Tools.FileIO;
8  import Framework.Rand;
9  import static Examples._6CompetitiveRelease.ExampleModel.*;
10 import static Framework.Util.*;
```

**1:** The package statement specifies where the file exists in the larger project structure

**2-7:** Imports all of the classes that we will need for the program.

**8:** Imports the static fields of the model so that we can use the type names defined there in the Agent class.

**9:** Imports the static functions of the Util file, which adds all of the Util functions to the current namespace, so we can natively call them. Statically importing Util is recommended for every project.

#### 10.9 Model Results

Table 3 displays the model visualization at timestep 0, timestep 400, timestep 1100, timestep 5500, and timestep 10,000 recorded from the GridWindow ToPNG function. The caption explores the notable trends visible in each image. Fig 4 displays the population sizes as recorded by the FileIO Write function at the end of every timestep.

| Timestep | No Drug / Constant Drug / Pulsed Drug |
|---|---|
| 0 | 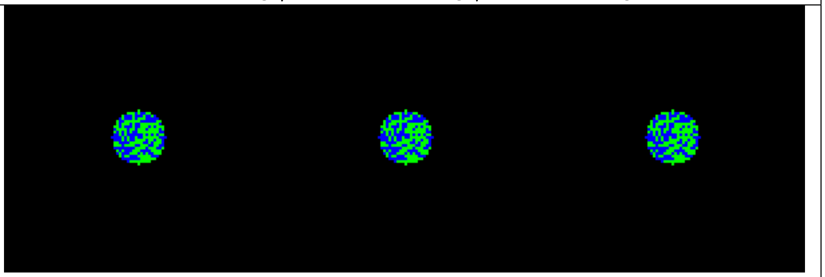 |
| 400 | 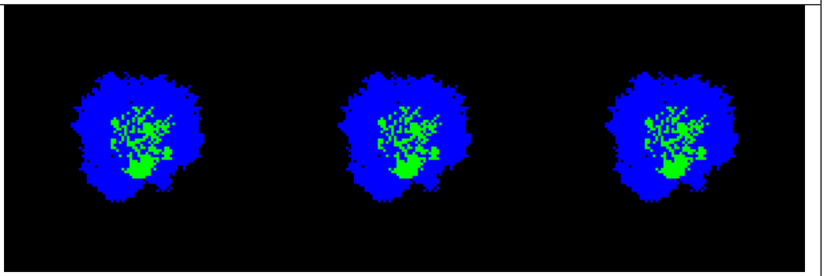 |
| 1100 | 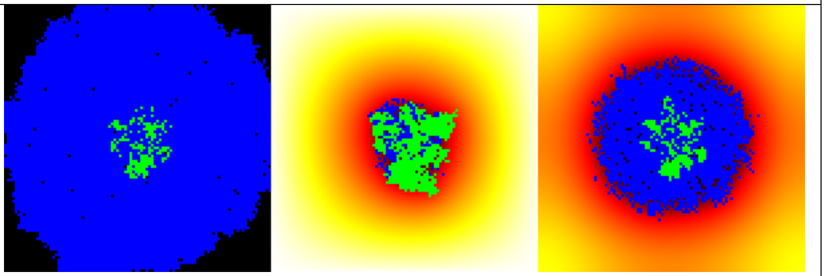 |
| 5500 | 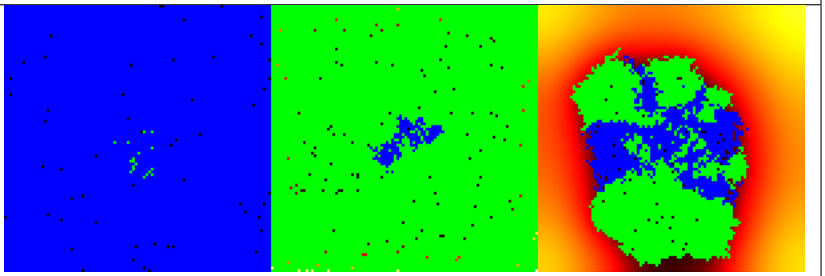 |
| 10000 | 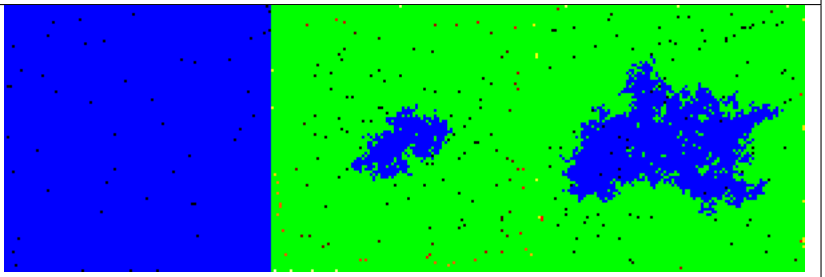 |

Table 3: Selected model visualization PNGs. Blue cells are drug sensitive, Green cells are drug resistant, background heatmap colors show drug concentration.

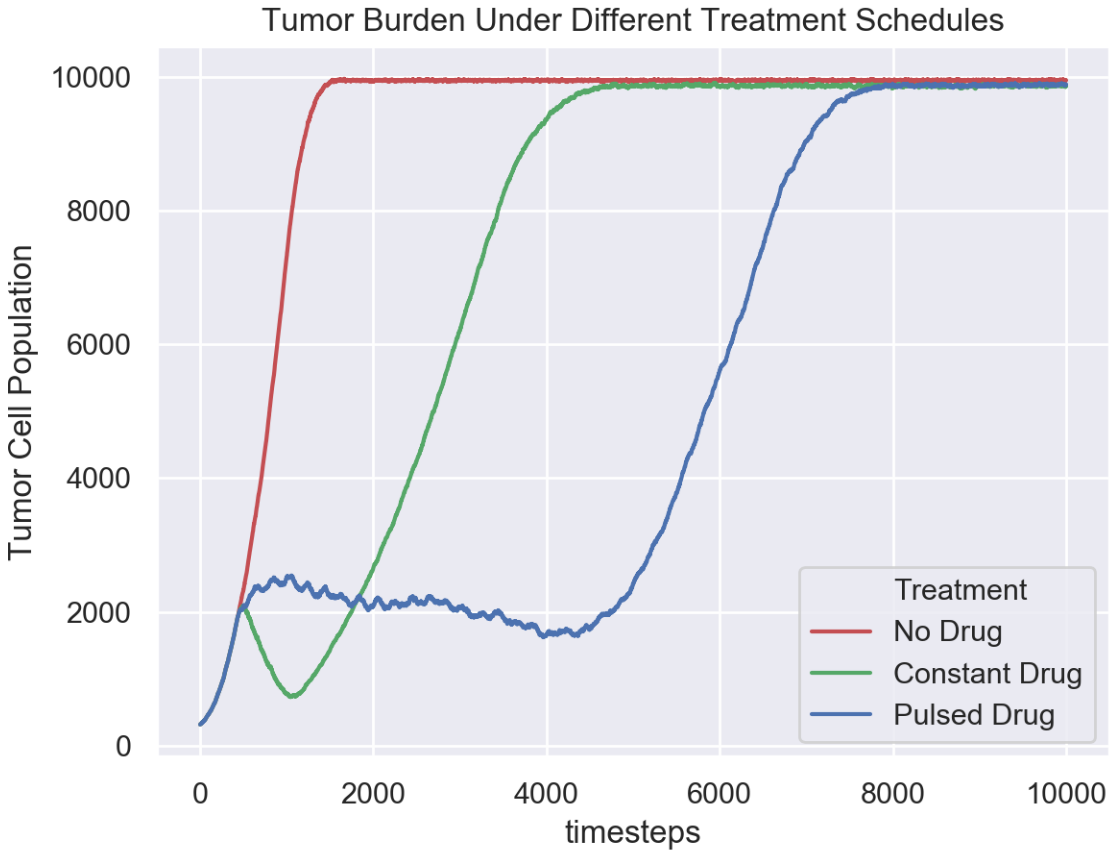

Figure 4: FileIO population output. This plot summarizes the changes in tumor burden over time for each model. This plot was constructed in python using data accumulated in the program output CSV file. Displayed using Seaborn with Python.

This example illustrates the power of HAL's approach to model building. Writing relatively little complex code, we setup a three model experiment with nontrivial dynamics along with methods to collect data and visualize the models. We now briefly review the model results.

As can be seen in Table 3, at timestep 0 and timestep 400 (right before drug application starts), all 3 models are identical. At timestep 1100 the differences in treatment application show different effects: when no drug is applied, the rapidly dividing sensitive cells quickly fill the domain, while when drug is applied constantly, the resistant cells overtake the sensitive population. Pulsed drug kills some sensitive cells, but leaves enough alive to prevent growth of the resistant cells. At timestep 5500, the resistant cells have begun to emerge from the center of the pulsed drug model. At timestep 10000, all domains are filled. Interestingly, in the models with drug application, the sensitive cells are able to survive in the center of the domain because drug is consumed by cells on the outside. This creates a drug-free zone in which the sensitive cells out-compete the resistant cells even when drug is applied constantly.

As can be seen in Fig 4, the pulsed therapy is the most effective at preventing tumor growth, however the resistant cells ultimately succeed in breaking out of the tumor center and out-competing the sensitive cells on the fringes of the tumor. It may be possible to contain a population of sensitive and resistant cells for longer by using a different pulsing schedule or by modifying the treatment schedule in response to the tumor growth (adaptive therapy). As the presented model is primarily an example, we do not explore how to improve treatment further. For a more detailed exploration of the potential of adaptive therapy for prolonging competitive release, see [1].

#### References

[1] Jill A Gallaher, Pedro M Enriquez-Navas, Kimberly A Luddy, Robert A Gatenby, and Alexander RA Anderson. Adaptive vs continuous cancer therapy: Exploiting space and trade-offs in drug scheduling.

#### Acknowelgements

This work was possible through the generous support of NIH funding, Anderson and Tessi acknowledge NCI U54CA193489, Anderson and Bravo acknowledge NCI UH2CA203781.
